# A weak link with actin organizes tight junctions to control epithelial permeability

**DOI:** 10.1101/805689

**Authors:** Brian Belardi, Tiama Hamkins-Indik, Andrew R. Harris, Daniel A. Fletcher

## Abstract

In vertebrates, epithelial permeability is regulated by the tight junction (TJ) formed by specialized adhesive membrane proteins, adaptor proteins, and the actin cytoskeleton. Despite the TJ’s critical physiological role, a molecular-level understanding of how TJ assembly sets the permeability of epithelial tissue is lacking. Here, we identify a 28-amino acid sequence in the TJ adaptor protein ZO-1 that is responsible for actin binding and show that this interaction is essential for TJ permeability. In contrast to the strong interactions at the adherens junction, we find that the affinity between ZO-1 and actin is surprisingly weak, and we propose a model based on kinetic trapping to explain how affinity could affect TJ assembly. Finally, by tuning the affinity of ZO-1 to actin, we demonstrate that epithelial monolayers can be engineered with a spectrum of permeabilities, which points to a new target for treating transport disorders and improving drug delivery.

## INTRODUCTION

In epithelial cells, the actin cytoskeleton plays numerous roles, ranging from structural to mechanical in nature. At the apical surface, cells make use of actin to assemble microvilli (Crawley et al., 2014), structures that increase the surface area of the membrane and expand the absorption potential of organs like the small intestine. On the basal side of polarized epithelia, focal adhesions rely on actin to orient and transmit forces between cells and extracellular matrix (Carragher and Frame, 2004). Actin filaments are also found at lateral contacts between epithelial cells, where they gird the cell in a belt-like arrangement (Miyoshi and Takai, 2008) and co-localize with two lateral junctions: the tight junction (TJ) and the adherens junction (AJ) (Fig. 1A).

**Figure 1.**
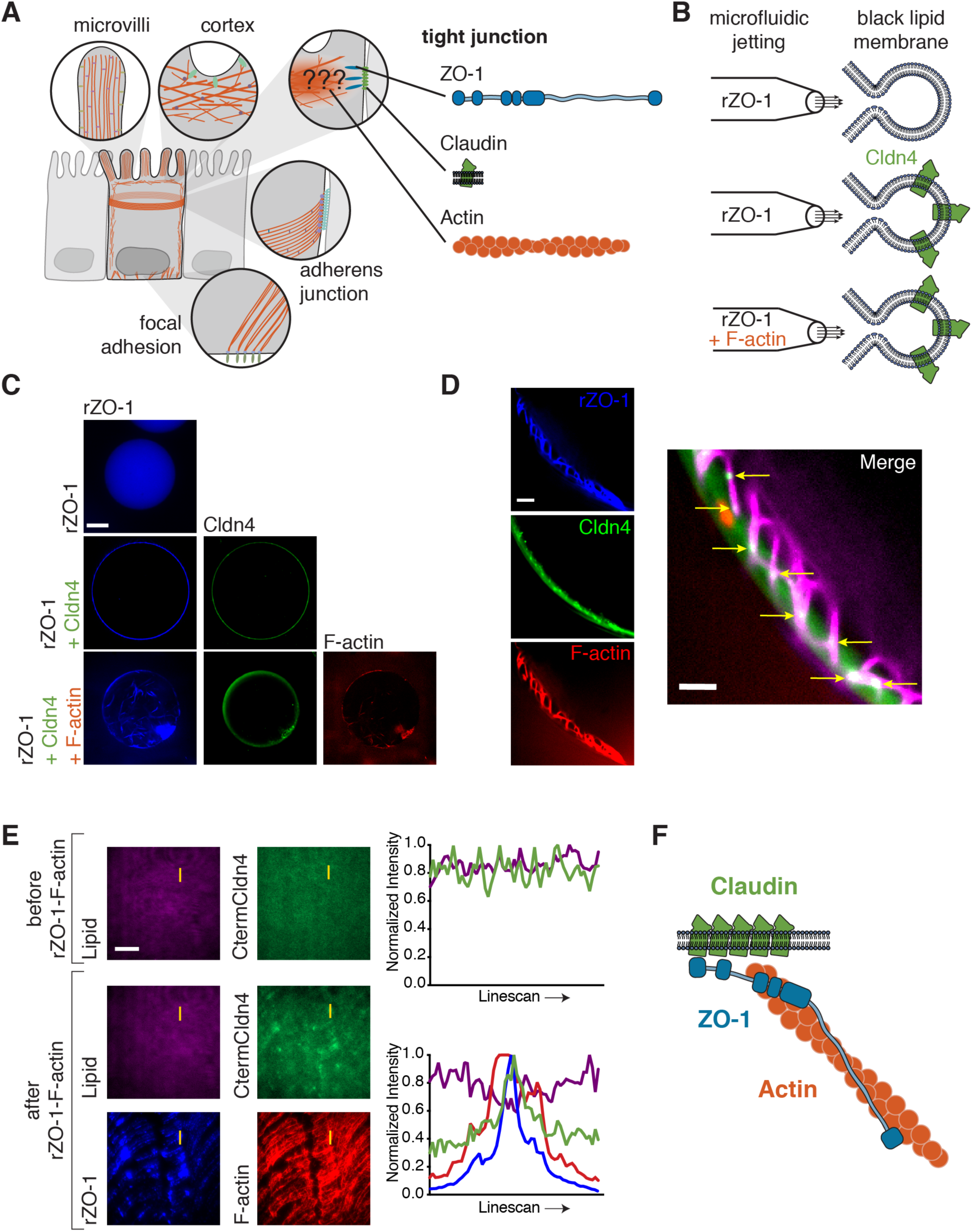
ZO-1 simultaneously engages actin and transmembrane claudins in vitro. (A) Schematic of actin structures in polarized epithelial cells. The role of actin at TJs is still unresolved. Depicted are three TJ components that lie in and near the membrane: the plaque protein, ZO-1, which possesses binding motifs for claudins and F-actin; transmembrane claudins; and filamentous actin. (B) GUVs were jetted with either rZO-1 (top), rZO-1 and Cldn4 (middle), or rZO-1, Cldn4, and F-actin (bottom). (C) Fluorescent micrographs of jetted GUVs under the three conditions outlined in (B). Scale bar, 50 μm. (D) Fluorescent micrographs of jetted GUVs containing rZO-1, Cldn4 and F-actin. Scale bar, 2.5 μm. The yellow arrows (right) indicate positions of enrichment of Cldn4 by ZO-1-F-actin meshes. (E) Fluorescent micrographs of CtermCldn4 peptide tethered to DOPC-based supported lipids bilayers in the presence or absence of ZO-1-F-actin complexes and line scan (yellow line). Scale bar, 20 μm. (F) ZO-1 simultaneously links the actin cytoskeleton with claudin in vitro.

The apical-most junction in vertebrate epithelia is the TJ, which is responsible for forming the vital barrier against paracellular flux, including at the blood-brain barrier and at the intestinal lining of the gut (Zihni et al., 2016). This is accomplished by selective pores generated by a class of adhesive membrane proteins known as the claudins (Furuse et al., 1998). In *trans*, claudins form extracellular channels between adjacent cells (Günzel and Yu, 2013; Suzuki et al., 2014), while in *cis*, claudins are thought to oligomerize to form polymeric strands within the same membrane (Gong et al., 2015; Irudayanathan et al., 2018; Koval, 2013; Piontek et al., 2007, 2011; Rossa et al., 2014; Sasaki et al., 2003; Zhao et al., 2018). In addition to the claudins, an abundant set of proteins including other membrane proteins, adaptors and actin have also been found to localize to the TJ in experiments based on fractionation, yeast two-hybrid screens, and proteomics (Mattagajasingh et al., 2000; Pulimeno et al., 2011; Van Itallie et al., 2013; Vogelmann and Nelson, 2005). Despite the extensive list of TJ-associated proteins, a mechanistic understanding of how key components assemble and interact with actin to establish a robust, yet malleable, paracellular barrier remains unresolved.

In contrast to TJs, a mechanistic understanding of actin’s role within the AJ, which neighbors the TJ, has emerged over the last two decades. Bundles of actin filaments stabilize the AJ and lend integrity to epithelial sheets (Yonemura, 2011). At the molecular level, actin is connected to the membrane at the AJ indirectly through a protein hierarchy that extends perpendicular to the lipid bilayer, beginning with membrane proteins, which are linked to cytoplasmic adaptors that are coupled to cytoskeletal proteins (Bertocchi et al., 2017). One critical complex that spans this space is that between the adhesive transmembrane protein E-cadherin, the cytoplasmic adaptors β- and α-catenin, and filamentous actin (F-actin) (Yamada et al., 2005). Early work on α-catenin pointed to the necessity of associating with actin, as its actin-binding domain (ABD) was required for maintaining durable contacts between cells (Nagafuchi et al., 1994). More recently, work from several groups has shown that the interaction between α-catenin and F-actin is of high affinity (K_D_ ∼ 400 nM) (Hansen et al., 2013) and is, notably, catch-bond-dependent (maximal lifetime at ∼8 pN of force) (Buckley et al., 2014). These experiments point to a tensile model of AJ assembly, where forces due to formin-directed actomyosin contractility (Acharya et al., 2017; Kobielak et al., 2004) and *trans* interactions across cells lead to strong association between the catenin complex and actin, thereby stabilizing the AJ and integrating the cortices of individual cells into a resilient mechanical continuum.

TJs are often compared to AJs because of their biochemical similarities (Hartsock and Nelson, 2008). Akin to E-cadherin, transmembrane claudin proteins have a cytoplasmic binding sequence, a PDZ-binding sequence at the C-termini, which links TJ membrane proteins to cytoplasmic adaptors that in turn bind to F-actin. The main adaptors at the TJ are the PDZ-domain-containing proteins ZO-1 and ZO-2, which are large, multi-domain proteins that possess actin-binding activity through an unidentified motif in their long, C-terminal disordered regions (Fanning et al., 2002). Despite mirroring molecular hierarchies, however, there are hints of significant differences between actin’s function at AJs and TJs. For instance, parallel bundles of actin filaments that are part of the AJs are absent from TJs (Hull and Staehelin, 1979). Similarly, actin-based contractility does not appear to play the same role at the two junctions. Inhibition of actomyosin contractility disrupts the AJ, significantly reducing the area of E-cadherin-based junctions (Liu et al., 2010) and the overall tissue stiffness of epithelial monolayers (Harris et al., 2014). Conversely, we and others have found that inhibiting actomyosin contractility, either by treatment with blebbistatin or with MLCK and ROCK inhibitors, improves the barrier function of the TJ (Supplemental Fig. 1) (Graham et al., 2019; Spadaro et al., 2017; Van Itallie et al., 2009; Yu et al., 2010). These experiments point to a fundamentally different role for actin at the TJ than for the neighboring AJ.

Here we describe how ZO-1 weakly couples the actin cytoskeleton to adhesive membrane proteins at the TJ to control the permeability of epithelial monolayers. This coupling is directed by a small, 28-amino acid actin-binding site (ABS) embedded in the middle of ZO-1’s C-terminal disordered region. We find that ZO-1’s ABS peptide, which represents a new actin-binding motif, is critical for TJ barrier function in epithelial cells. In contrast to the AJ, we show that low affinity association between ZO-1 and the actin cytoskeleton is critical for assembling TJs with robust barrier function, and we propose a kinetic trap model to explain the different role the actin cytoskeleton plays at TJs compared to AJs. By modulating the affinity of ZO-1 to F-actin, we were able to generate a series of epithelial cell lines with both increased and decreased permeabilities. Our data suggest that ZO-1’s interaction with actin could be a new therapeutic target for efforts to address both delivering drugs past the blood-brain barrier (BBB) and for treating transport disorders, such as inflammatory bowel disease.

## RESULTS

### Reconstitution of ZO-1 with claudin and F-actin forms co-localized structures in vitro

Previous work has found that three TJ proteins – membrane claudins, the ZO adaptor proteins and filamentous actin (F-actin) – interact and are indispensable for barrier function (Fanning et al., 1998; Itoh et al., 1999) (Fig. 1A). First, we sought to verify the binding of these proteins in vitro and observe how they self-organize, similar to how in vitro binding assays of AJ proteins showed how the catenins bound to the C-terminus of E-cadherin and to F-actin (Drees et al., 2005). To visualize complex formation at a membrane surface, we relied on a method we previously developed to embed the four-pass transmembrane claudins into the lipid bilayers of giant unilamellar vesicles (GUVs) (Belardi et al., 2019). This method is based on microfluidic jetting of black lipid membranes and gives rise to GUVs containing oriented claudin proteins, wherein claudin’s C-terminal PDZ binding motif faces the lumen of vesicles. We took advantage of the microfluidic jetting procedure to load into the lumen of giant vesicles a recombinant, truncated form of ZO-1 that contains claudin- and F-actin-binding domains and regions (rZO-1, see Materials and Methods for details) (Fig. 1B). We found that rZO-1 localized exclusively to the claudin-4 (Cldn4)-containing membrane, whereas in the absence of Cldn4, rZO-1 was strictly lumenal (Fig. 1C).

Next, we loaded both rZO-1 and F-actin into the lumen of Cldn4-containing giant vesicles. In these vesicles, we observed that rZO-1 and F-actin formed an interconnected structure that appeared to template the enrichment of Cldn4 along the lipid bilayer at sites of co-localization with F-actin and rZO-1 (Fig. 1C and 1D). To further verify three-component complex formation, we turned to supported lipid bilayers (SLBs) and total internal reflection fluorescence (TIRF) microscopy. As a surrogate for full-length Cldn4, we conjugated the C-terminal sequence of Cldn4, which contains the PDZ binding motif, to maleimide phospholipids in the SLB (Lin et al., 2014). After reaction, the Cldn4 peptide was imaged by TIRF and appeared homogenous on the membrane (Fig. 1E, top). To the Cldn4 peptide SLBs, we next added the rZO-1-F-actin mixture. Similar to the GUV results above, we observed a heterogeneous distribution after incubation (Fig. 1E, bottom), where enrichment of the Cldn4 peptide co-localized with rZO-1-F-actin meshes. We performed fluorescence recovery after photobleaching (FRAP) to determine if the mobility of the Cldn4 peptide was influenced by the presence of rZO-1 and F-actin, and we found that, indeed, its mobility was reduced and that the Cldn4 peptide’s organization was templated by rZO-1-F-actin tracks (Supplemental Fig. 2). Taken together, these results suggest that ZO-1 is capable of bridging F-actin and claudins simultaneously, forming co-localized structures (Fig. 1F).

### ZO-1 binds to F-actin through a 28 amino acid motif within its C-terminal disordered region

How exactly human ZO-1 interacts with F-actin has remained unclear, beyond that it requires a segment in ZO-1’s C-terminal disordered region (Fanning et al., 2002). So, we next turned our attention to identifying the specific site responsible for F-actin binding (Fig. 2A). To do this, we developed a simple, cell-free protein expression assay to rapidly screen sequences within ZO-1 for F-actin binding. In this assay, plasmids containing ZO-1 sequences are combined with cell extracts to initiate transcription and translation. These mixtures are then placed on a glass surface in order to immobilize expressed ZO-1 sequences. To this mixture, phalloidin-stabilized F-actin is added and fluorescence microscopy is used to visualize F-actin binding at the surface (Fig. 2B and 2C). We started with a segment of the C-terminal disordered region, known as the actin-binding region (ABR), to identify the critical residues for F-actin binding. A modified binary search was used, examining the N-terminal, C-terminal, or middle half of ZO-1 stretches. Doing so iteratively, we identified a 28-amino acid sequence within the ABR, which we refer to as the actin-binding site (ABS), that is the minimal sequence within ZO-1 capable of F-actin binding (Fig. 2D). In the context of ZO-1’s primary sequence, the ABS resides in the central portion of the ABR of ZO-1 and, interestingly, has no homology to other actin-binding proteins in humans.

**Figure 2.**
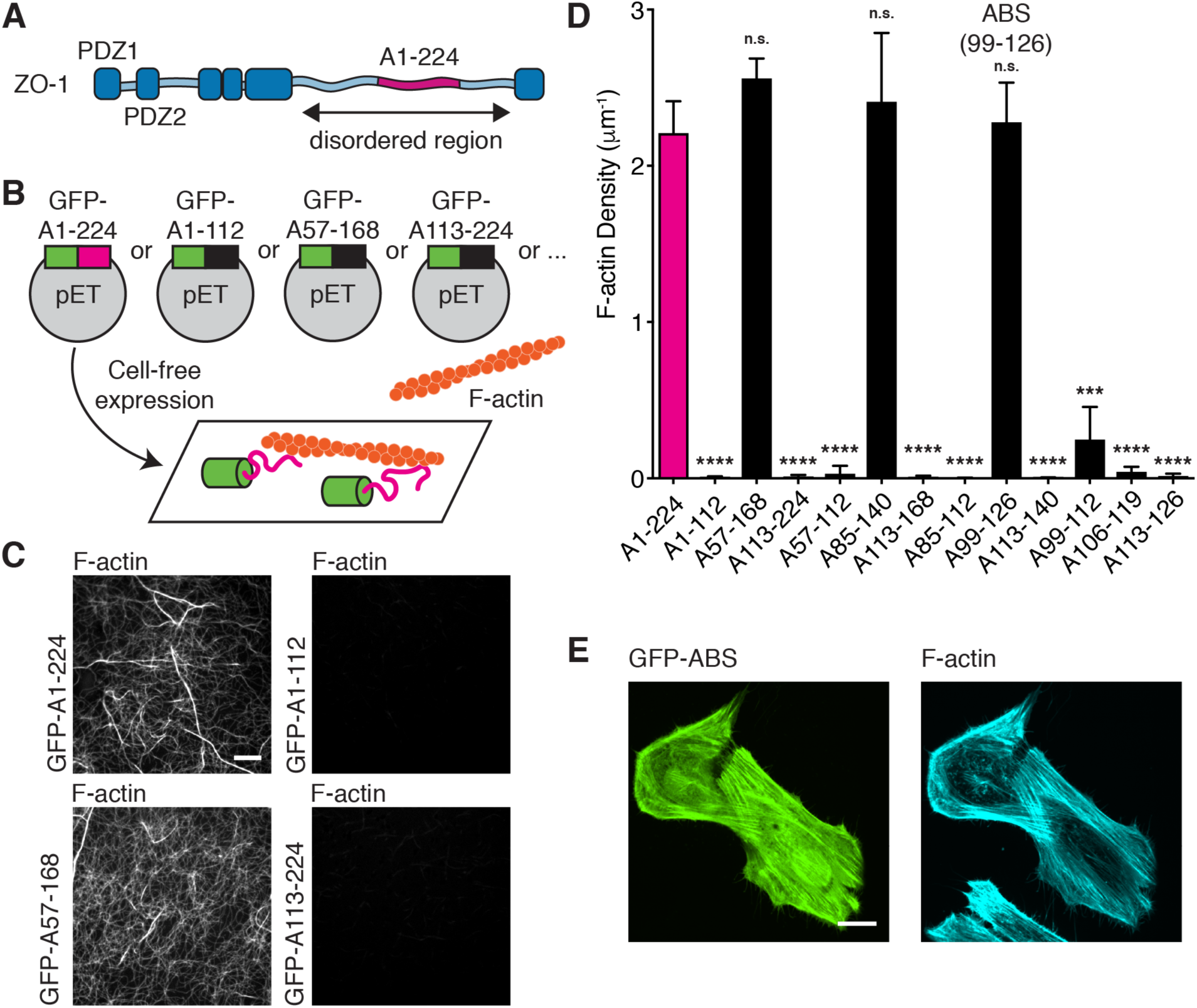
Identification of an ABS within ZO-1’s C-terminal disordered region. (A) Schematic of human ZO-1’s primary sequence. A search for ZO-1’s ABS was performed on a 224 amino acid portion (magenta) of ZO-1’s long C-terminal disordered region. (B) Schematic of cell-free, actin-binding assay. Segments of ZO-1 were fused to GFP and expressed using cell-free expression. F-actin binding was visualized using fluorescence microscopy. (C) Fluorescent micrographs of F-actin binding assay after expressing different segments of ZO-1’s disordered region. Scale bar, 10 μm. (D) Quantification of F-actin density from cell-free actin-binding assay for various constructs encoding different portions of the disordered region. The magenta bar highlights the full 224 amino acid region, while the yellow bar represents the minimal ABS of ZO-1. Bars represent mean ± SD, n=3, (p-values determined using a two-sample t-test with A1-224, *** p<0.001, **** p<0.0001, n.s. p>0.05). (E) Fluorescent micrographs of HeLa cells expressing GFP-ABS (left). Cells were fixed and stained for F-actin using AF647-phalloidin (right). Scale bar, 20 μm.

We performed an alanine scan of the ABS to identify amino acids that were indispensable for F-actin binding. Unsurprisingly, most of the 28 amino acids were important for F-actin binding, especially the positively-charged residues (Supplemental Fig. 3A). However, two alanine mutant sequences, M1 and M6, had low and comparable binding of F-actin, respectively. To verify the binding interaction in cells, we expressed a fluorescently-labeled minimal ABS (GFP-ABS) in HeLa cells, which provide distinct actin structures that can be used to characterize actin-binding domains (Harris et al., 2019). In this context, we found that the ABS decorated actin filaments in stress fiber structures as well as lamellipodia (Fig. 2E). To explore evolutionary conservation of the ABS, we aligned the site across ZO-1 homologs in various metazoans (Supplemental Fig. 3B). We noted that vertebrates appeared to retain the site with a high degree of identity, whereas the ABS sequence is absent from aligned ABR regions in invertebrate ZO-1 homologs.

### The 28-amino acid Actin Binding Site (ABS) of ZO-1 is required for barrier function

Our in vitro studies led us to ask whether ZO’s ABS is necessary for physiological functions of the TJ, for instance the barrier function of epithelial cells. If so, then impaired barrier function should be observable after deletion of the ABS from ZO proteins. Using CRISPR/Cas9, we generated knock out cell lines of either ZO-1, ZO-2, or both ZO-1 and ZO-2 (dKO) in MDCK II cells (see Materials and Methods for details). By immunostaining, we confirmed knockouts of the ZO proteins and found that proper localization of other TJ proteins, for example claudins, occludin, and cingulin, was disrupted in the dKO line but not in single KO cells (Fig. 3A and Supplemental Fig. 4A and 4B) (Otani et al., 2019). We next measured barrier function of the different ZO knockouts by transepithelial electrical resistance (TEER) and a fluorescence Transwell permeability assay. In both assays, we observed that barrier function was severely compromised in the dKO line but not in wildtype or in the single KO cells (Fig. 3B and Supplemental Fig. 4E).

**Figure 3.**
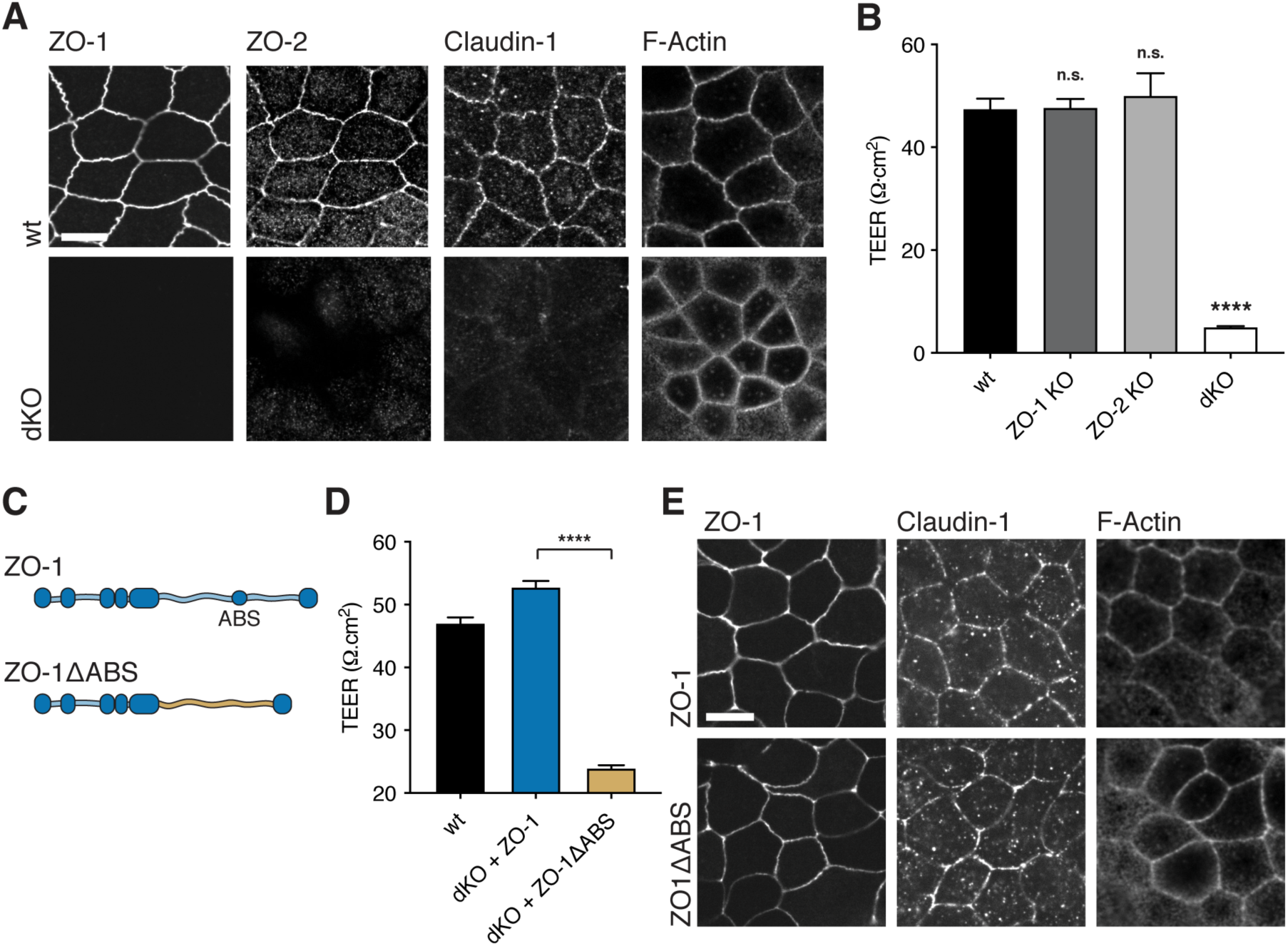
ZO-1’s ABS is necessary for robust barrier function in dKO MDCK cells. (A) Immunofluorescent micrographs of TJ proteins in wt and dKO cells. The images show lack of ZO-1 and ZO-2 in dKO cells. The localization of TJ proteins, e.g. Claudin-1 and F-actin, are altered in dKO cells. Scale bar, 10 μm. (B) TEER measurements of wildtype (wt) MDCK II cells and of CRISPR/Cas9-generated ZO-1 and ZO-2 KO cells at day 4 of confluency. Bars represent mean ± SEM (p-values determined using a multi-comparison ANOVA between each of the means, **** p<0.0001, n.s. p>0.05). N values represent biological replicates, wt (N=6), ZO-1 KO (N=4), ZO-2 KO (N=3), dKO (N=7). Barrier function was abolished for cells lacking ZO-1 and ZO-2 (dKO). (C) Schematic of ZO-1 constructs with and without ZO-1’s ABS introduced into dKO cells. (D) TEER measurements of wt and dKO cells expressing full-length ZO-1 and ZO-1ΔABS. Measurements are from day 4 of confluency. Bars represent mean ± SEM, n=4, (p-values determined using a two-sample t-test, **** p<0.0001). (E) Immunofluorescent micrographs of ZO-1, Claudin-1, and F-actin in dKO cells expressing ZO-1 and ZO-1ΔABS. No difference in localization of TJ proteins was observed between the two cell lines. Scale bar, 10 μm.

Next, we turned to the question of whether the 28-amino acid ABS contributed to ZO-1’s ability to establish a paracellular barrier in epithelial cells. To test this, we first expressed full-length human ZO-1 in dKO cells. In this case, barrier function was fully restored upon ZO-1 expression (Fig. 3D). We then tested a ZO-1 construct that lacked the ABS (ΔABS) in the dKO. In the case of ΔABS, barrier function was significantly impaired compared to the construct that possessed the native ABS (Fig. 3D). Both proteins appeared to traffic to the TJ normally by confocal microscopy, and immunostaining also confirmed proper protein localization and density of other TJ proteins in both cell lines (Fig. 3E and Supplemental Fig. 4D). These studies suggest that ZO-1 and F-actin, in concert, influence the establishment of an epithelial TJ with robust barrier function.

### Replacing the ABS of ZO-1 with actin network regulators does not rescue barrier function

Having shown the importance of ZO-1’s ABS on epithelial barrier function, we next asked what role the ABS might be playing in the organization of actin networks at the TJ. Several Rho family GTPases and their regulatory effectors, guanine nucleotide exchange factors (GEFs) and GTPase activating proteins (GAPs), localize to the TJ, including Cdc42, RhoA (Quiros and Nusrat, 2014), p114RhoGEF (Terry et al., 2011), GEF H1 (Aijaz et al., 2005), Tuba (Otani et al., 2006), among others, which implies that specific actin structure and activity are important for assembling TJs with robust barrier function. In fact, Hansen et al. showed that α-catenin’s ABD directly inhibits branched actin architectures at AJs (Hansen et al., 2013). This led us to wonder whether the ABS might be partly responsible for influencing the actin architecture at the TJ in epithelial cells.

To test this, we replaced the ABS of ZO-1 with GEFs known to act on different Rho GTPases (Fig. 4A), inducing specific actin structures and activity. Specifically, we inserted the catalytic domains, DH or DH/PH domains, from three well-characterized GEFs – LARG, intersectin-1, and Tiam1 – into ZO-1. The GEF from LARG has previously been characterized to act on the GTPase RhoA and has been used ectopically to generate parallel actin bundles and contractility (Wagner and Glotzer, 2016), while the GEFs from intersectin-1 (ISTN) and Tiam1 act on Cdc42 and Rac1, respectively, which have also been used ectopically to engineer branched actin structures in cells (Beco et al., 2018; Zimmerman et al., 2017). By confocal microscopy, all three constructs localized to the TJ in the dKO cell line (Fig. 4C). We observed slightly flatter TJs for the LARG construct, whereas both the ISTN and Tiam1 constructs generated TJs that were more tortuous than wildtype – consistent with the known activities of the three GEFs. We next used TEER to evaluate the effect of GEF insertion on barrier function. All three constructs failed to rescue barrier function in dKO cells compared to full-length ZO-1 (Fig. 4B), although each case achieved slightly elevated resistance values compared to the construct lacking an inserted GEF. Interestingly, both ISTN and Tiam1 constructs achieved higher TEER than the LARG construct, which might indicate a preference for branched actin structures at the TJ.

**Figure 4.**
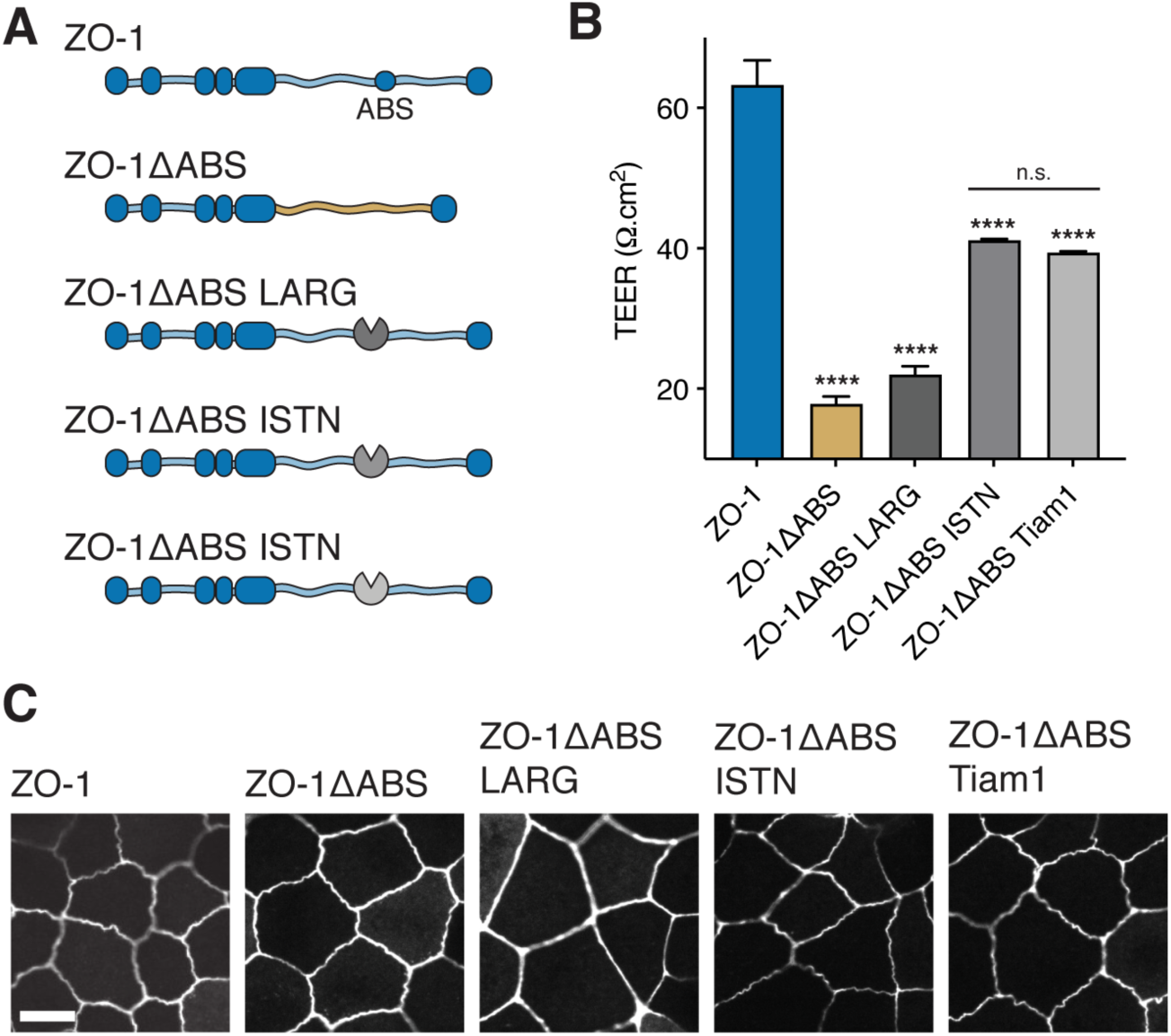
Engineering actin network structures at the TJ fails to restore barrier function of cells lacking ZO-1’s ABS. (A) Schematic of ZO-1 constructs introduced into dKO cells with the actin regulators, LARG, ISTN, and Tiam1, replacing the ABS. (B) TEER measurements of dKO cells expressing ZO-1ΔABS GEF constructs. Measurements are from day 4 of confluency. Bars represent mean ± SEM, n=3, (p-values determined using a multiple comparison one-way ANOVA, p-values represent comparison with ZO-1, **** p<0.0001, n.s. p>0.05). (C) Fluorescent micrographs of ZO-1ΔABS GEF constructs show localization to the TJ membrane. Scale bar, 10 μm.

### Barrier function is reduced when the ZO-1 ABS is replaced with a high-affinity actin-binding domain

Since insertion of the LARG, ISTN, and Tiam1 GEFs into ZO-1 lacked the ability to directly bind F-actin, we reasoned that the direct interface between ZO-1 and F-actin must be necessary for establishing a robust barrier in epithelial monolayers. If true, then replacing ZO-1’s ABS with another actin-binding domain (ABD) should restore barrier function. To test this, we replaced ZO-1’s ABS with an ABD from another junctional protein, α-catenin (Fig. 5A). As described above, α-catenin is part of the AJ and provides necessary contacts between the E-cadherin/catenin complex and F-actin. After confirming that the ZO-1 α-Catenin construct localized to the TJ (Fig. 5B), we measured TEER to assay for barrier function. Interestingly, the construct did not rescue barrier function compared to full-length ZO-1 (Fig. 5B), despite a slight improvement in resistance compared to the ZO-1 construct lacking ABS (ΔABS).

**Figure 5.**
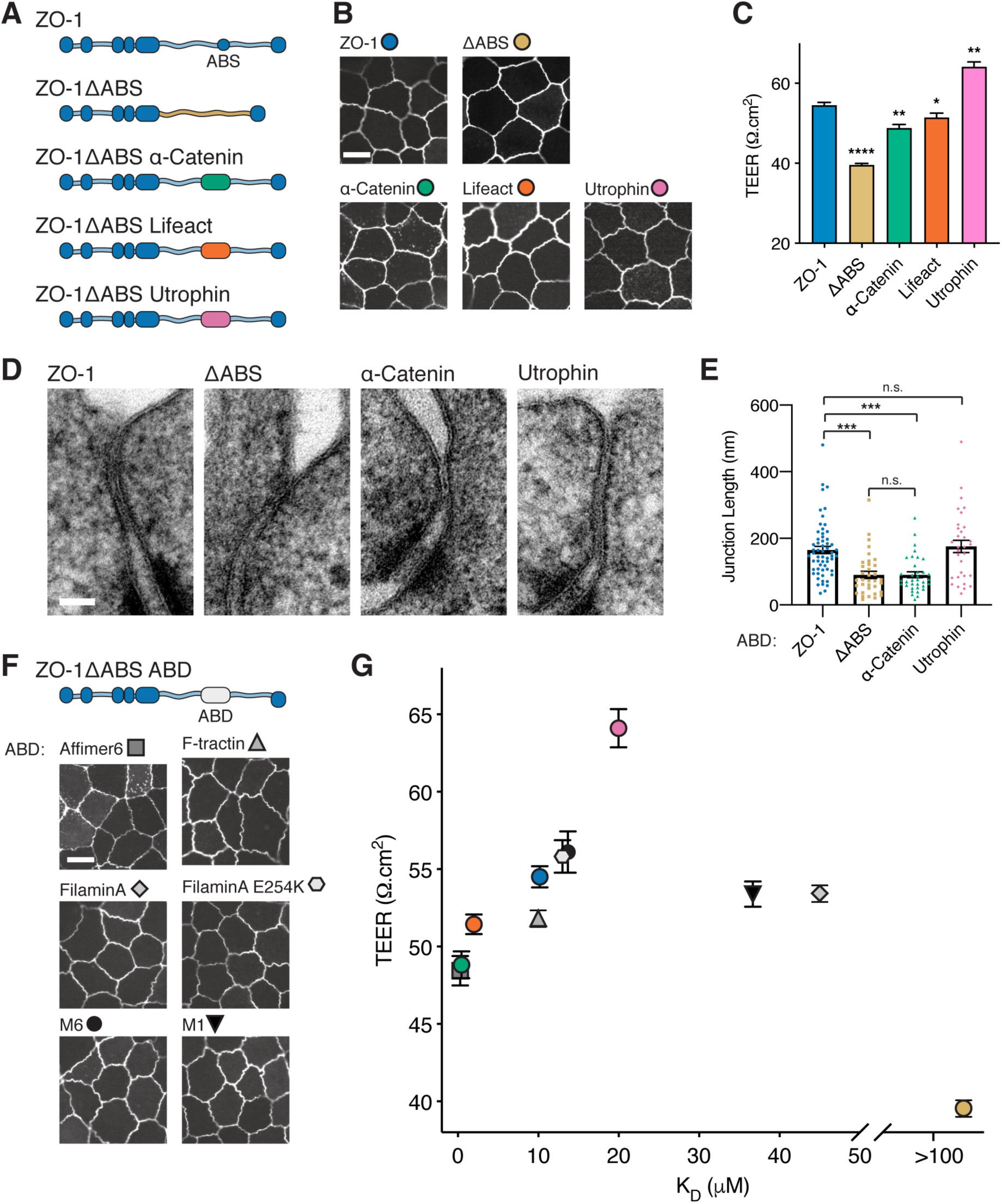
Weak association between ZO-1 and F-actin is required for robust epithelial barrier function. (A) Schematic of ZO-1 constructs introduced into dKO cells with ectopic ABDs replacing the native ABS of ZO-1. (B) Fluorescent micrographs of dKO cells from day 4 of confluency expressing ZO-1ΔABS with the ABDs from the AJ protein, α-Catenin, the short peptide, Lifeact, and the tandem calponin-homology domain protein, Utrophin. All constructs localized to the TJ. Scale bar, 10 μm. (C) TEER measurements of dKO cells expressing ZO-1ΔABS with the ABDs from α-Catenin, Lifeact, and Utrophin. Measurements are from day 4 of confluency. Bars represent mean ± SEM, n=3, (p-values determined using a two-sample t-test for ZO-1 vs. ABD, * p<0.05, ** p<0.01, **** p<0.0001). (D) Representative TEM images of dKO cells expressing ZO-1 constructs with various ABDs. Scale bar, 50 nm. (E) Quantification of TJ length from TEM images of dKO cells expressing ZO-1 constructs with various ABDs. Bars represent mean ± SEM, (p-values determined using a multi-comparison one-way ANOVA with a Brown-Forsythe test, *** p<0.001, n.s. p>0.05). Each symbol represents one cell-cell junction, ZO-1 (n=59), ΔABS (n=35), α-Catenin (n=35), Utrophin (n=32). (F) Schematic of ZO-1 constructs introduced into dKO cells with ectopic ABDs replacing the native ABS of ZO-1 (top). Fluorescent micrographs of ZO-1ΔABS ABD constructs in dKO cells (bottom). All constructs localized to the TJ. Scale bar, 10 μm. (G) TEER measurements of dKO cells expressing ZO-1ΔABS with ectopic ABDs replacing ZO-1’s ABS plotted vs. the affinity of the ABD for F-actin (see Table S2). Measurements are from day 4 of confluency. Symbols represent mean ± SEM, n=3.

The ABD from α-catenin is a 236-amino acid, five-helix bundle, which is a significant deviation in size and fold from ZO-1’s ABS that could disrupt the ZO-1-F-actin interface. To address this concern, we replaced the ABS with an ABD of similar size, namely the short peptide Lifeact, which consists of 17 amino acids (Riedl et al., 2008). However, the ZO-1 Lifeact construct also failed to fully recover barrier function (Fig. 5E), although we noted an improvement in barrier function over the α-Catenin construct. Besides size and fold, another parameter that might have an effect on the proper assembly of TJ complexes is the affinity between binding partners.

### Barrier function is rescued and can be enhanced when the ZO-1 ABS is replaced with low-affinity actin-binding domains

Both α-Catenin and Lifeact are examples of ABDs with high affinity (low K_D_) toward F-actin. We measured the K_D_ for ZO-1’s ABS and found that its affinity for F-actin was much weaker (∼10 μM for both ABS and ABR, Supplemental Fig. 5). To test whether weak association of ZO-1 with F-actin was an important feature of the TJ, we decided to replace the ABS with an ABD that has a weak affinity for F-actin. Utrophin, a calponin homology (CH) domain-containing protein, has an ABD with a K_D_ of ∼20 μM toward F-actin (Winder et al., 1995). After installing Utrophin’s ABD in ZO-1 and measuring TEER of an epithelial monolayer formed with the construct, we found that the ZO-1 containing the Utrophin ABD not only completely recovered TEER compared to full-length ZO-1 but that the cell line expressing ZO-1 Utrophin led to enhanced barrier function in relation to wildtype ZO-1 (Fig. 5B). To visualize sub-cellular physical differences between the cell lines expressing different forms of ZO-1, we used transmission electron microscopy (TEM) to image the TJ length. Compared to wt ZO-1, we found that cells expressing either ZO-1 lacking actin-binding (ΔABS) or ZO-1 α-Catenin had shorter junctional lengths (Fig. 5C and 5D). We also found that wt ZO-1 and ZO-1 Utrophin, constructs with weak association to F-actin, possessed long junctional lengths (Fig. 5C and 5D). Longer TJs may provide more opportunities for claudin-claudin contacts across cells, therefore leading to the lower permeability of monolayers expressing wildtype ZO-1 or ZO-1 Utrophin.

Intrigued by this finding, we replaced the ABS of ZO-1 with an affinity series of other ABDs in order to better characterize the F-actin affinity-barrier function relationship for ZO-1. With this approach, we found that the relationship between ZO-1’s affinity for F-actin and the ultimate permeability of the epithelial monolayer was defined by a bell-shaped curve, with a maximum at low affinity (∼20 μM) and reduced barrier function at high and very low affinities (Fig. 5E and Table S2). In each case, proper localization of ZO-1 constructs was observed (Fig. 5F). Taken together, these studies show that the claudin-ZO-1-F-actin interface is necessary for achieving robust barrier function in epithelial monolayers. However, in contrast to the AJ where high affinity is important for junction integrity, these data demonstrate that a weak association between ZO-1 and F-actin is critical for assembling proper barrier function in epithelial cells. This insight enabled us to engineer epithelial monolayers with either diminished or enhanced barrier function by modulating the affinity of ZO-1 for F-actin.

### Simulations show that low-affinity interactions between two polymers prevent kinetic trapping in misaligned configurations

How might weak association to F-actin be leveraged by TJs to assemble robust barrier function? One possibility might stem from a unique feature of claudins, which is their ability to form polymeric structures, or strands, in the plane of the membrane in cells (Gong et al., 2015; Irudayanathan et al., 2018; Koval, 2013; Piontek et al., 2007, 2011; Rossa et al., 2014; Sasaki et al., 2003; Zhao et al., 2018). These structures differ from transmembrane protein dimers or protein clusters in that each claudin monomer contains a front-to-back contact that repeats to generate a linear polymeric chain (Suzuki et al., 2014). Using structured illumination microscopy, recent work by van Itallie et al. noted a surprising arrangement of ectopically expressed claudin polymers in fibroblasts (Van Itallie et al., 2016). They found that in the presence of ZO-1, claudin polymers appeared to align with actin filaments, whereas in the absence of ZO-1, claudin polymers had no directional correlation with actin filaments. This observation led us to wonder whether the dynamics of aligning one polymer with another might be influenced by their interfacial affinity.

To examine this, we simulated fluctuating polymers (green) on a 2-D lattice, where static polymers (orange) were arranged in a grid pattern (Fig. 6A). Affinity between the two polymers was varied, and alignment of the two-polymer system was monitored over time. Figure 6B shows representative traces of alignment vs. time for a single fluctuating polymer with varying affinities. Under low affinity conditions, the fluctuating polymer reached a steady-state alignment in a short period of time (∼100 τ, where τ represents the time step) and oscillated around this alignment for the rest of the simulation. By contrast, under high affinity conditions, the fluctuating polymer remained in low alignment configurations for extended period of times (>10,000 τ) as the simulation evolved to higher alignments. We next compiled simulations from 100 different starting configurations. We found that increasing the affinity by an order of magnitude resulted in an average dwell time difference of over an order of magnitude (Fig. 6C), suggesting long periods of stalling for high affinity polymers before alignment is maximized. Figure 6D plots the normalized average alignment at different numbers of time steps as a function of affinity. After 100 τ, the lowest affinity polymer had already reached ∼97% of its equilibrium alignment, while the highest affinity had only reached ∼65% of its steady-state alignment.

**Figure 6.**
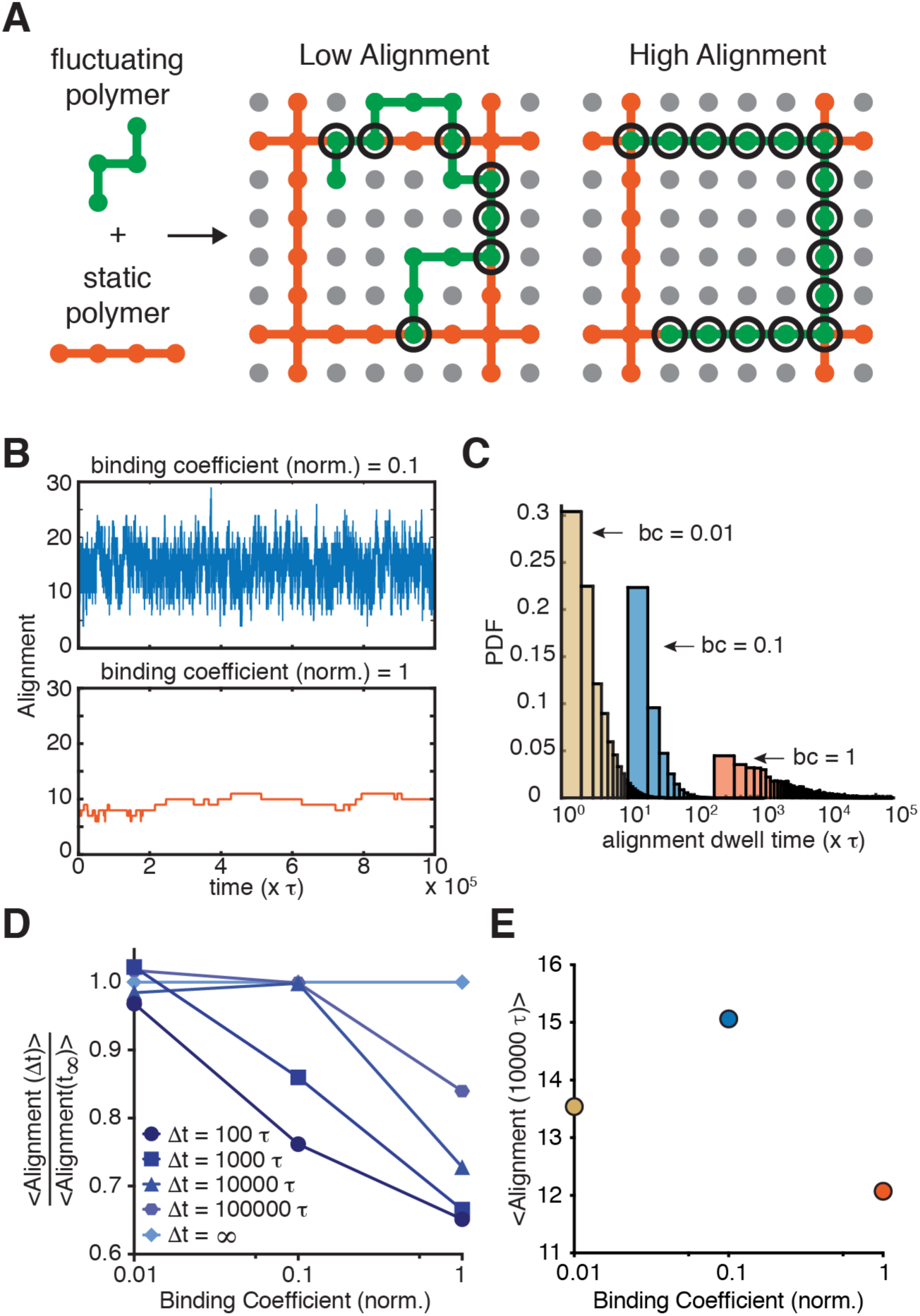
High affinity interaction between two polymeric species gives rise to kinetic traps. (A) Schematic of lattice model indicating low and high alignment between a static polymer (orange) and a fluctuating polymer (green). (B) Alignment traces of a single polymer fluctuating under two binding coefficients over time. (C) Probability density function of alignment dwell times of fluctuating polymers under various binding coefficients (bc). Dwell time refers to the number of steps that polymers remain in individual alignment configurations. (D) Normalized average alignments of fluctuating polymers under various binding coefficients at different time intervals. At short time intervals, fluctuating polymers with large binding coefficients are trapped at low alignments, suggesting that a high affinity interaction to F-actin at the TJ may kinetically trap claudins in unfavorable configurations. (E) Average alignment of fluctuation polymers under various binding coefficients at 10,000τ. At this time interval, kinetic trapping is convolved with the total number of contacts between the two polymers, resulting in a bell-shaped curve.

Collectively, these data point to a key feature of a two-polymer system – that the high affinity interaction of one polymer with another polymer will kinetically trap the system in sub-optimal configurations. Considering also that affinity affects the maximum level of alignment that can be achieved, Figure 6E presents non-normalized alignment after 10,000 τ showing that very low affinity, despite not being kinetically trapped, fails to reach high alignment, revealing a bell-shaped curve reminiscent of what we see experimentally. These results suggest that weak interactions between ZO-1 and F-actin may be critical for allowing claudin polymers to achieve the alignment and organization necessary for achieving robust barrier function Other factors, such as variations in fluctuating polymer length and flexibility, as well as differences in linear polymer density, geometry, and turnover, would be expected to alter the magnitude of alignment predicted by this model, providing the cell with additional ways to use kinetic trapping to tune TJ barrier function.

## DISCUSSION

The TJ and AJ are neighbors along the lateral surface of epithelial cells, and together have been hypothesized to work as a single sub-cellular structure called the apical junction complex (AJC) (Roignot et al., 2013). While a detailed understanding of TJ interactions with actin has remained incomplete, a picture of actin’s function at the AJ has now come into focus (Bertocchi et al., 2017). Parallel actin bundles are present at the AJ (Hull and Staehelin, 1979), and actin-based contractility is critical for the strong lateral adhesions that maintain integrity across epithelial tissue (Harris et al., 2014).

Our work shows that the actin cytoskeleton functions fundamentally differently at the TJ. Actin-based contractility (Supplemental Fig. 1) appears detrimental to the TJ and its associated barrier function. More significantly, a weak, not strong, linkage between the actin cytoskeleton and claudins drives TJ organization and assembly – a stark contrast to the high-affinity F-actin association at AJs (Hansen et al., 2013). This distinction extends to the molecular structures that interface with F-actin. We find that TJ assembly is directed by a small peptide embedded in the long, C-terminal disordered region of ZO-1, while the same interfaces at AJs are defined by an ABD with a five-helix fold. We also show that although actin association is critical at TJs, replacing ZO-1’s F-actin with α-catenin affinity diminishes barrier function in epithelial monolayers. Taken together, our work suggests that the role for actin across epithelial junctions is not uniform.

How, then, might α-catenin’s and other high-affinity actin-binding domains drive sub-optimal arrangement of TJs? Our Monte Carlo simulations provide one possible explanation. We found that high affinity interactions between a two-polymer system leads to kinetic trapping, i.e. long-lived local minima rather than a global minimum along an energetic landscape. Kinetic traps in biology are not unprecedented. In fact, kinetic traps (in metastable states) have been directly observed in S-layer protein assembly (Shin et al., 2012) using atomic force microscopy and during the folding of proteins, including insulin (Hua et al., 1995). Thus, it’s possible that the ZO-1 α-Catenin construct forms a kinetically trapped structure at the TJ. Our results suggest that the non-equilibrium dynamics of complex assembly, as opposed to equilibrium-based phenomena, might be critical for organizing the TJ, much as they have recently been shown to be important for carboxysome assembly (Rotskoff and Geissler, 2018). We imagine that TJs are also taking advantage of low-affinity interactions and dynamics to arrive at configurations that give rise to robust but malleable barrier function in epithelium.

Since the ABS represents a short stretch within a much longer sequence of ZO-1’s disordered region, we wondered how changes to the disordered region might affect barrier function. To obtain an initial perspective on this, we examined post-translational modifications of ZO-1 as barrier function is established in monolayers. One modification previously examined in the literature is ZO-1 phosphorylation, which was suggested to change upon TJ formation (Howarth et al., 1994; Stevenson et al., 1989). Based on this evidence, we used proteomics to compare phosphorylation of ZO-1’s C-terminal disordered region, which contains the ABS, before and after the establishment of TJs. We observed extensive modification within ZO-1’s disordered region – a total of 32 different sites of phosphorylation (Supplemental Fig. 6A). Comparing the number of phosphorylated ZO-1 sites within this region before and after TJ formation, we found that the extent of phosphorylation of ZO-1’s disordered region increased by a factor of three after the TJ was established (Supplemental Fig. 6B).

How might these phosphorylation events within the disordered region affect barrier function? To address this, we created three mutant forms of ZO-1: a phosphomimetic version (PhosAll ZO-1) with aspartic acid residues at each of the 32 sites of phosphorylation, a different phosphomimetic version with aspartic acid residues at modified serine and threonine residues (PhosSerThr ZO-1), and a phospho-null mutant (PhosNull-ZO-1) with either alanine or phenylalanine residues at each of the 32 sites (Supplemental Fig. 6C). After introducing these three constructs into dKO cells, we measured TEER for each cell line and found that the final TEER values for the PhosNull-ZO-1 was higher than either of the phosphomimetic ZO-1 constructs (Supplemental Fig. 6D). These data indicate that modifications to ZO-1’s disordered region do indeed affect the barrier function of TJs, possibly by changing accessibility of the ABS. Furthermore, the data point to the fact that the disordered region of ZO-1 is dynamically altered to adjust epithelial permeability – an indication that multiple states of ABS-directed TJ organization are accessible to wildtype cells.

In conclusion, our findings suggest that a major function of F-actin in epithelial cells is to organize and stabilize claudin strands in the membrane through weak interactions with the adaptor protein ZO-1. In this capacity, F-actin at TJs acts not to apply strong forces to the junction, which is the case at AJs, but to template and align the transmembrane proteins responsible for forming intermolecular pores across epithelial cells. Our work also suggests that actin’s templating role at the TJ could offer a new therapeutic target. We found that by modifying the affinity of ZO-1 to F-actin, the permeability of epithelial monolayers could be manipulated to exhibit either diminished or enhanced barrier function. The bioavailability of small molecule and protein therapeutics would benefit greatly from the ability to selectively modulate the paracellular flux between epithelial cells. By temporarily increasing the affinity of ZO-1 for F-actin or completely inhibiting F-actin association, drug delivery past epithelial monolayers, through the intestinal walls of the gut or through the BBB, could be improved. Moreover, patients with transport disorders characterized by leaky epithelium, for instance inflammatory bowel diseases, need treatments that specifically restore barrier function (Turner, 2009). This could also be accomplished by varying the affinity of ZO-1 to F-actin. One possible advantage of this strategy is that targeting the ZO-1-F-actin interface does not abolish barrier function. Consequently, a drug modulating ZO-1-F-actin association might lead to less toxicity and less side-effects than other general permeability enhancers, such as ultrasound treatment (Bors and Erdő, 2019) and sodium caprate administration (McCartney et al., 2016). Our work points to altering ZO-F-actin interactions as an exciting area of future investigation with the promise of fine-tuning barrier properties in the gut or at the BBB as a means therapeutic intervention.

## MATERIALS AND METHODS

### General methods

All of the chemical reagents were of analytical grade, obtained from commercial suppliers, and used without further purification, unless otherwise noted. Alexa Fluor 647 phalloidin was purchased from ThermoFisher. 1,2-diphytanoyl-*sn*-glycero-3-phophoscholine (DPhPC), 1,2-dioleoyl-*sn*-glycero-3-phosphocholine (DOPC), 1,2-dioleoyl-*sn*-glycero-3-phosphoethanolamine-N-[methoxy(polyethylene glycol)-2000], ammonium salt (DOPE-PEG), 1,2-dioleoyl-*sn*-glycero-3-phosphoethanolamine-N-[4-(p-maleimidophenyl)butyramide], sodium salt (DOPE-MPB) were obtained from Avanti Polar Lipids, Inc. 1,2-Dioleoyl-*sn*-glycero-3-phosphoethanolamine labeled with Atto 390 (DOPE-Atto 390) and Lucifer yellow were purchased from Atto-tec and Sigma Aldrich, respectively.

Fluorescence imaging was carried out on a Ti Eclipse microscope (Nikon) equipped with a CSU-X spinning disk confocal module (Yokogawa) and a Zyla sCMOS camera (Andor). Fluorescence micrographs of giant vesicles or cells were acquired with either a 20x objective (Nikon, NA 0.45) or a 60x objective (Nikon, NA 1.49 TIRF). TIRF imaging was performed on the Ti Eclipse microscope (Nikon) using a 60x objective (Nikon, NA 1.49 TIRF) and an iXon Ultra EM-CCD camera (Andor).

### Protein expression and purification

ZO-1 proteins used for in vitro reconstitution and co-sedimentation experiments were all prepared using insect cell expression. The rZO-1 construct consisted of an N-terminal RFP tag followed by amino acids 1-411 of the human ZO-1 gene, containing PDZ1 and PDZ2 domains, in frame with amino acids 1159-1382 of the human ZO-1 gene, followed by the Dual Strep purification tag. For co-sedimentation assays, ZO-1 ABR constructs consisted of an N-terminal EGFP tag followed by amino acids 1159-1382 of the human ZO-1 gene and a C-terminal Dual Strep purification tag. GFP-ABRM1 and GFP-ABRM6 constructs contained a stretch of four alanine mutations at different positions of the ABS sequence, amino acids 1-4 and 21-24 of the ABS, respectively. The ABS construct consisted of an N-terminal GST solubilization tag and thrombin cleavage site followed by amino acids 1257-1284 of the human ZO-1 gene (ABS) in frame with EGFP and a C-terminal 6xHis tag. For bacmid production, each sequence was cloned into the pFastBac HTA vector and transformed into DH10Bac bacterial cells. Transformed cells were grown on LB agar plates with kanamycin (50 μg/mL), gentamycin (7 μg/mL), tetracycline (10 μg/mL), IPTG (40 μg/mL), and Bluo-gal (100 μg/mL) and colonies that were white in color were picked for amplification and isolation of bacmid. PCR of bacmids confirmed the insertion of ZO-1 sequences. Sf-9 cells were transfected with isolated bacmids using Cellfectin (ThermoFisher), and after 4 days, supernatants containing baculovirus were collected and clarified by centrifugation at 1,000 x g for 5 min. Virus was then amplified by two rounds of Sf-9 cell infection and supernatant collection.

For expression, concentrated baculovirus was used to infect Sf-9 in a 1 L culture at 27 °C. After 2 days, cells were collected by centrifugation at 400 x g for 10 min and lysed into a lysis buffer containing 25 mM HEPES, pH 7.5, 150 mM NaCl, 1 mM EDTA, 1 mM TCEP supplemented with DNase I and protease inhibitors using a Dounce homogenizer. The lysate was clarified by centrifugation at 18,000 rpm at 4 °C. ZO-1 constructs were then purified using affinity chromatography. Briefly, clarified lysate was cycled over a Strep column (IBA Lifesciences) for 2 hr at 4 °C. The column was then washed with lysis buffer. For elution, lysis buffer containing 2.5 mM desthiobiotin was added to the column, and eluted proteins were further purified by size exclusion chromatography using a Superdex 75 column (GE Healthcare) into a buffer containing 25 mM HEPES, pH 7.5, 150 mM NaCl, 1 mM TCEP.

### Microfluidic jetting of giant unilamellar vesicles (GUVs)

GUVs were formed by placing a microfluidic jetting nozzle (Microfab Technologies, Inc., single jet microdispensing device with 25 μm orifice) filled with a 0.2 μM rZO-1, 10% OptiPrep (Sigma-Aldrich), 25 mM HEPES, pH 7.5, 150 mM NaCl solution in close proximity, <200 μm, to DPhPC planar bilayers with and without embedded GFP-Cldn4 (see Belardi et al., 2019). The rZO-1-containing solution ultimately constitutes the lumen of the jetted GUVs and has a matched osmolarity, but different density, compared to the outer buffer. Jetting was performed with and without AF647-phalloidin-stabilized F-actin (0.4 μM) included in the rZO-1-containing solution. The piezoelectric actuator of the nozzle was controlled by a waveform generator (Agilent) and an amplifier (Krohn-Hite) with pulse train envelops designed by a custom Matlab script. For jetting of GUVs, the actuator was triggered by an increasing parabolic envelop defined by 40 trapezoidal bursts at 15 kHz, 3 µs rise and fall times, a 30 µs hold time, and a maximum voltage of 15-30 V. Planar bilayer deformation and GUV formation were monitored using brightfield microscopy with a high-speed camera (Photron, 1024PCI). GUVs formed by microfluidic jetting sunk to the poly-L-lysine-coated coverglass due to the density mismatch between the interior of the GUVs and the surrounding buffer and were imaged using spinning disk confocal microscopy.

### CtermCldn4 supported lipid bilayers

CtermCldn4-functionalized supported lipid bilayers (SLBs) were prepared in a similar manner to Lin et al. (Lin et al., 2014). Briefly, glass coverslips were RCA cleaned and treated with a 1 mg/mL solution of SUVs containing DOPC (97%), DOPE-MPB (2.5%), DOPE-PEG (1%) and DOPE-Atto 390 (0.1%) for 10 min to form SLBs. SLBs were washed five times with a volume of 200 μL of buffer containing 25 mM HEPES, pH 7.5, 150 mM NaCl. After washing, SLBs were blocked by treating with β-casein (Sigma Aldrich) for 10 min at room temperature. SLBs were washed again three times with a volume of 200 μL of buffer containing 25 mM HEPES, pH 7.5, 150 mM NaCl. DOPE-MPB lipids were then reacted with the terminal cysteine residue of a fluorescent CtermCldn4 peptide (300 nM, 5-FAM-Ahx-CPPRTDKPYSAKYSAARSAAASNYV, GenScript) for 1 hr at room temperature. Unreacted maleimide lipids were capped by treating SLBs with 2-mercaptoethanol for 10 min. SLBs were washed five times with a volume of 200 μL of buffer containing 25 mM HEPES, pH 7.5, 150 mM NaCl. Imaging of SLBs was performed using TIRF microscopy with a 60x objective before addition of rZO-1 and F-actin. AF647-phalloidin-stabilized F-actin (0.4 μM) was combined with rZO-1 (0.2 uM) for 30 min before addition to SLBs. rZO-1-F-actin complexes were incubated with SLBs for 30 min at room temperature and then washed three times with a volume of 200 μL of buffer containing 25 mM HEPES, pH 7.5, 150 mM NaCl. SLBs were again imaged using TIRF microscopy. Fluorescence recovery after photobleaching (FRAP) of fluorescent CtermCldn4 was performed by narrowing a field stop in the excitation path and illuminating the sample at high power for 10 s. The field stop was then opened and a time-lapse acquisition (every 20 s) was performed.

### Cell free expression and in vitro actin-binding assay

To express a range of GFP-tagged ZO-1 segments, cell-free protein expression was performed by first cloning appropriate sequences of ZO-1 into a pET28a expression vector. Expression of GFP fusions of ZO-1 segments was commenced by combining 3.5 uL of bacterial extract (Biotech Rabbit), 4 uL of reaction buffer (Biotech Rabbit), 2 uL of plasmid solution (final mass, 1 ug of DNA), and 0.5 uL of a 20 mM IPTG solution and placing the mixture at 37 °C for 1 hr. A laser-cut imaging chamber (acrylic wells UV-cured to 1.5 glass coverslip) was cleaned by sonicating the glass in the presence of deionized water, EtOH, and 3M NaOH solutions, sequentially. After washing with deionized water, the cell-free expression mixture was applied to chambers to immobilize the expressed fluorescent constructs. To this mixture, AF647-phalloidin-stabilized F-actin was added to a final concentration of 0.5 μM. After 45 min, each chamber was imaged using fluorescence microscopy without washing.

### ABS alignment

To align the ABS site across species, ZO-1 homologs were identified by a protein BLAST (NCBI) search: human (Uniprot Q07157), mouse (Uniprot P39447), rat (Uniprot A0A0G2K2P5), dog (Uniprot O97758), frog (Uniprot A0A1L8GT63), zebrafish (Uniprot A0A2R8QMK9), chicken (Uniprot A0A1D5NWX9), fruit fly (Uniprot A0A0B4K6Y7), roundworm (Uniprot Q8I103), sea urchin (Uniprot W4ZFF8), sea squirt (Uniprot A0A3Q0JQ32), lancelet (Uniprot C3YKF2), acorn worm (NCBI ProteinID XP_006813237.1), brachiopod (Uniprot A0A2R2MJX0), pacific oyster (NCBI XP_011435800.2), leech (Uniprot T1FAR7), flatworm (Uniprot G4VGU1), hydra (Uniprot Q9BKL2), and placozoan (Uniprot B3S7T9). No ZO-1 homologs were found in the ctenophore and the porifera clades. A local alignment of the human ZO-1 ABR was performed (Water, EMBL) for each ZO-1 homolog. With the homologous ABR sequences in hand, we then performed a second local alignment (Water, EMBL) with the 28-amino acid ABS sequence.

### Co-sedimentation actin-binding assay

F-actin was prepared by polymerizing β-actin at 68 µM for 1.5 hr at room temperature. Various concentrations of F-actin were then combined with a constant concentration of GFP-tagged ZO-1 constructs (0.5 μM) in Buffer F. Sub-stoichiometric concentrations of ZO-1 constructs were used in all experiments, such that the assumption of [F-actin]_total_ ≈ [F-actin]_free_ was valid. After incubation at room temperature for 30 min, F-actin was pelleted at 150,000 x *g* for 60 min at 4 °C. The supernatants were then collected, and unbound ZO-1 construct fluorescence intensity was analyzed using a fluorimeter (Biotek Instruments, Inc.). Bound fractions were fitted with the following equation 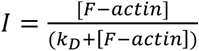, where *I* is the bound fraction, [F-actin] is the F-actin concentration and *k_D_* is the dissociation constant.

### Cell culture and cell lines

MDCK II cells were a gift from Keith Mostov (UCSF) and maintained at 37°C and 5% CO_2_ in high glucose DMEM (4.5 g/l), supplemented with 10% fetal bovine serum (FBS) and penicillin-streptomycin (pen-strep). ZO-1 constructs were cloned using PCR and Gibson assembly into pHR backbone plasmids. Cell lines were created through lentivirus infection. Briefly, HEK293 cells were transfected with TransIT-293 (Mirus) according to manufacturer’s instructions with three plasmids, pMD.2g, p8.91 and pHR with ZO-1 constructs (see Table for amino acids details). Cells were grown for 2 days, after which media was collected and virus was concentrated with Lenti-X (Clontech) according to manufacturer’s instructions. Virus was added to freshly passaged MDCK II cells, and cells were grown for two days before passaging and removing media. Cell lines created with fluorescently tagged proteins were sorted and normalized for expression using the UC Berkeley Flow Cytometry Facility (BD Bioscience Influx Sorter). Cell lines were confirmed with confocal imaging and immunoblot.

ZO-1, ZO-2, and double knockout (dKO) cell lines were created by first building a Cas9-expressing MDCK II cell line. Briefly, a lentiCas9-Blast (Addgene) plasmid was used to create stably expressing-Cas9 cell lines by selection under 5 ug/mL blasticidin for 7 days. Cas9 expression was confirmed with immunoblot. Three guide RNAs (gRNAs) for TJP1 were examined to create the ZO-1 KO (Table S1). The three pLenti-gRNA-puro plasmids were transduced into Cas9-expressing cells before selection in 10 ug/mL puromycin for 6 days. Knockout was initially confirmed with immunofluorescence imaging of ZO-1. Clonal cell lines were created by dilution plating, where a single cell suspension (5 cells/mL) was plated in a 96-well tissue culture dish. On day 7, each well was checked for a single colony, and on day 14, cells were passaged. Knockout was verified with immunofluorescence, immunoblot, and genomic sequencing (Supplemental Figure 4). dKO cells were created by following the protocol above with multiple TJP2 gRNAs (Table S1) and several TJP1 KO clones. After double knockout verification, a clone generated from gRNA 2 for TJP1 and gRNA 1 for TJP2 was used in subsequent experiments (Table S1).

### Barrier assays

Cells were plated on 24-well Transwell inserts (polyester (PE), 0.4 μm pore size (Corning)) coated with 30 μg/mL Collagen I (Cellmatrix) at a cell density of 3.33E4 cells/cm^2^. Transepithelial electrical resistance measurements (TEER) was performed using the ENDOHM6 cup chamber (WPI) with EVOM2 (WPI) in cell culture media. For transport measurements, phenol red-free media (4.5 g/L glucose, (-) L-glut, (-) Sodium Pyruvate, (-) phenol red DMEM supplemented with 10% FBS, pen-strep, and GlutaMax) with 10 uM of Lucifer yellow was added to Transwell insert. After 3 hours of incubation, media was collected from the basal culture well and fluorescence was measured using a fluorescence plate reader (Biotek Instruments, Inc.). Apparent permeability (P_app_) was calculated as 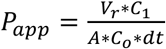, where V_r_ is the volume of the basal well, C_1_ is the concentration measured in the basal well, A is the area of the transwell insert, C_o_ is the concentration of Lucifer yellow added to the insert, and dt is the incubation time.

### Immunostaining and imaging

Cells were plated on a 2.1 ug/mL collagen I gel (Cellmatrix), pH 7.4, in a glass-bottom cell culture dish at a cell density of 3.33E4 cells/cm^2^ and grown for 2 or 4 days. Fixation methods were optimized for each antibody used. Cells were either fixed in 1% of 4% PFA at room temperature for 20 minutes or in 100% EtOH or 100% MeOH at −20° C for 30 min. Cells fixed in PFA were first permeabilized with 0.2% (v/v) Triton X-100 at room temperature for 10 minutes. To block, fixed cells were incubated in 5% (w/v) BSA in PBS at room temperature for 2 hours. Cells were then incubated with primary antibodies in a 1% (w/v) BSA in PBS solution at 4° C, overnight. Secondary antibodies were applied in a 1% BSA in PBS solution at room temperature for 1 hour. Three PBS washes were done after each antibody incubation step. Cells were incubated in Hoechst for 10 minutes at room temperature. Cells were imaged on a Ti Eclipse microscope (NIKON) using a 60x 1.49 NA objective and an iXon Ultra EMCCD (Andor). For live cell imaging, cells were plated identically, but not fixed, and imaged on the same microscope.

### Immunoblot

Cells were plated in a 6-well plate at a cell density of 3.33E4 cells/cm^2^ and grown for 2 or 4 days. Cells were lysed in RIPA buffer (1% (v/v) Triton X-100, 0.1% (w/v) SDS, 1% (w/v) deoxycholate) with protease inhibitors (Thermo Fisher) on ice for 30 minutes. Lysate was spun at 13,000 rpm for 15 minutes at 4° C. A BCA assay (Thermo Fisher) was performed according to manufacturer’s instructions to determine protein concentration. All samples were diluted to the same protein concentration. Loading buffer was added to lysate, and samples were run on a 4-20% polyacrylamide gel (Bio-Rad) and at 200 V for 35 minutes. Gel was transferred using iBlot (Thermo Fisher) according to manufacturer’s instructions at 20 V for 5 min. Ponceau S was incubated for 10 minutes at room temperature to visualize total protein. The blot was blocked in 2% (w/v) BSA in PBS with 0.1% Tween-20 (PBS-T) for 1 hour at room temperature. Primary antibody was administered in 1% (w/v) BSA overnight on a shaker at 4° C, and secondary was administered in 1% (w/v) BSA for 1 hour on a shaker at room temperature. Three, 10-minute PBS-T washes were performed after each antibody incubation. The blot was visualized on a ChemiDoc (BioRad).

### PCR of genomic DNA

Control and MDCK II cell lines were plated in a 6-well plate and grown to confluency. Genomic DNA was isolated using PureLink® Genomic DNA Mini Kit (Thermo Fisher) according to manufacturer’s instructions. PCR of ZO-1 was run on an agarose gel to confirm a single band, and the amplified segment was sequenced by the UC Berkeley DNA Sequencing Facility. PCR and sequencing were used to confirm knockout cell lines as well as insertion of ZO-1 and other constructs.

### Monte Carlo polymer simulations

A custom Matlab (Mathworks) script was developed to run dynamic simulations of 2-D polymeric species on a 2-D lattice. Polymer dynamics were modeled using the slithering snake algorithm (Wall and Mandel, 1975), which is well-suited to studying polymer fluctuations in crowded environments, e.g. cell membranes (Itzhak et al., 2016). Briefly, linear, static polymers were placed on a 100×100 lattice every 5 lattice sites apart in both the x and y directions. A second polymeric species with an odd-number of monomers was placed on the gird in a random configuration. A set of randomly orientated polymers with a constant number of contacts with the static polymers was created. The randomly oriented polymeric species was then initialized to perform a slithering snake movement. Either end of the polymer was chosen at random as the head of the polymer and allowed to sample adjacent lattice sites by comparing the transition probability to a random number. If a move is accepted, the head moves to the new lattice site and the rest of polymer follows. The transition probability depends on the number of contact sites and the effective affinity between the fluctuating polymer and the linear, static polymer. If the move position is already occupied, the head and tail of the polymer are switched, and the same procedure is repeated. We performed simulations of 100 initial starting conditions at three different polymer-polymer affinities. Data are displayed for polymer lengths of 39 monomers. The results are consistent for larger and smaller fluctuating polymers, however at lengths much smaller than the bounding static polymer squares, kinetic trapping is negligible.

### TEM

For transmission electron microscopy (TEM), cells were plated on 24-well Transwell inserts (polyester (PE), 0.4 μm pore size (Corning)) coated with 30 ng/mL Collagen I (Cellmatrix) at a cell density of 3.33E4 cells/cm^2^ and grown for 4 days. Cells were fixed for 1.5 hours at room temperature in 2.5% glutaraldehyde and paraformaldehyde (EMS) in 0.1 M sodium cacodylate buffer, pH 7.2, and then washed three times with sodium cacodylate buffer for 15 min. After fixation, membranes were cut out of Transwell inserts for handling and kept hydrated. Membranes with cells were then incubated in 1% osmium tetroxide in sodium cacodylate buffer for 30 minutes on ice in the dark, followed by three washes in sodium cacodylate buffer for 15 min. Samples were dehydrated using progressively higher percentages of ice cold EtOH (35%, 50%, 70%, 80%, 90%, 100%, 100%), each incubation was performed for 7 minutes. For TEM, after dehydration in EtOH, cells were infiltrated with resin with progressively higher percentages of resin (25%, 50%, 75%, 100%, 100%), each incubation was performed for 15 minutes. Samples were cut and layered into molds and cured at 60° C for two days. Sections, 70-90 nm thin, were cut on either a Reichert-Jung Ultracut E (Leica) or Leica EM UC6 (Leica) and collected onto 50 mesh copper grids, then post-stained with 2% aqueous uranyl acetate and lead citrate for 5-7 minutes in each. Images were collected on an FEI Tecnai12 transmission electron microscope (FEI, Hillsboro, OR).

### Phosphoproteomics of ZO-1

To identify phosphorylated residues, phosphoproteomics of ZO-1 was carried out on dKO MDCK II cells expressing wildtype ZO-1. Cells were cultured for either 6 hr or 4 days and then washed twice with PBS (1x). Lysis buffer containing 50 mM Tris, pH 7.5, 150 mM NaCl, 0.5% NP-40, 1% Triton X-100, 1 mM EDTA supplemented with protease and phosphatase inhibitors was added to cell monolayers for 5 min on ice. After scraping, cells in lysis buffer were incubated for 45 min at 4 °C with end-over-end mixing. Lysates were then clarified by centrifugation at 14,000 x g for 20 min. Supernatant was recovered and incubated with 40 μL of washed GFP Trap magnetic beads (ChromoTek) overnight at 4 °C according to the manufacturer’s instructions. Beads were washed five times with buffer containing 25 mM HEPES, pH 7.5, 150 mM NaCl, and 1 mM EDTA. Sample buffer was then added to the beads, and SDS-PAGE was performed on immunoprecipitated ZO-1. Protein was visualized using Coomassie staining and ZO-1 bands were excised from acrylamide gels. In-gel digestion with trypsin was then carried out, and recovered peptides were submitted for phosphoproteomic analysis through the QB3 Vincent J. Coates Proteomics facility.

## AUTHOR CONTRIBUTIONS

Conceptualization, B.B., T.H-I., D.A.F.; Methodology, B.B., T.H-I., and A.R.H.; Investigation, B.B. and T.H-I.; Formal Analysis, B.B. and T.H-I.; Software, B.B.; Writing – Original Draft, B.B., T.H-I., and D.A.F.; Writing – Review & Editing, B.B., T.H-I., A.R.H., and D.A.F.; Funding Acquisition, D.A.F., B.B., and A.R.H.; Resources, D.A.F.; Supervision, D.A.F.

## ACKNOWLEDGEMENTS

We thank Sho Takatori and Jordi Silvestre-Ryan for advice and helpful discussions and Prof. Jianghui Hou and his laboratory (Washington University School of Medicine in St. Louis) for the generous gift of GFP-Cldn4. We also thank Eva Schmid for critical reading of the manuscript, and the staff, especially Danielle Jorgens and Reena Zalpuri, at the University of California, Berkeley Electron Microscope Laboratory for advice and assistance in electron microscopy sample preparation and data collection. We acknowledge the assistance and support of the University of California, Berkeley Cell Culture Facility. This work was supported by grants from the NIH (R01GM114344), and the UCSF NSF Center for Cellular Construction (DBI-1548297). This work used the Vincent J. Proteomics/Mass Spectrometry Laboratory at UC Berkeley, supported in part by NIH S10 Instrumentation Grant S10RR025622. B.B. was supported by the NIH Ruth L. Kirschstein NRSA fellowship from the NIH (1F32GM115091). T.H-I was supported by an NSF-GRFP fellowship, Berkeley Stem Cell Center’s NIH Stem Cell Biological Engineering Training Program (T32GM098218), and as a UC Berkeley Lloyd Scholar. A.R.H. was in receipt of an EMBO long-term fellowship 1075–2013 and an HFSP fellowship LT000712/2014. D.A.F. is a Chan Zuckerberg Biohub Investigator.

---

**Supplemental Figure 1.**
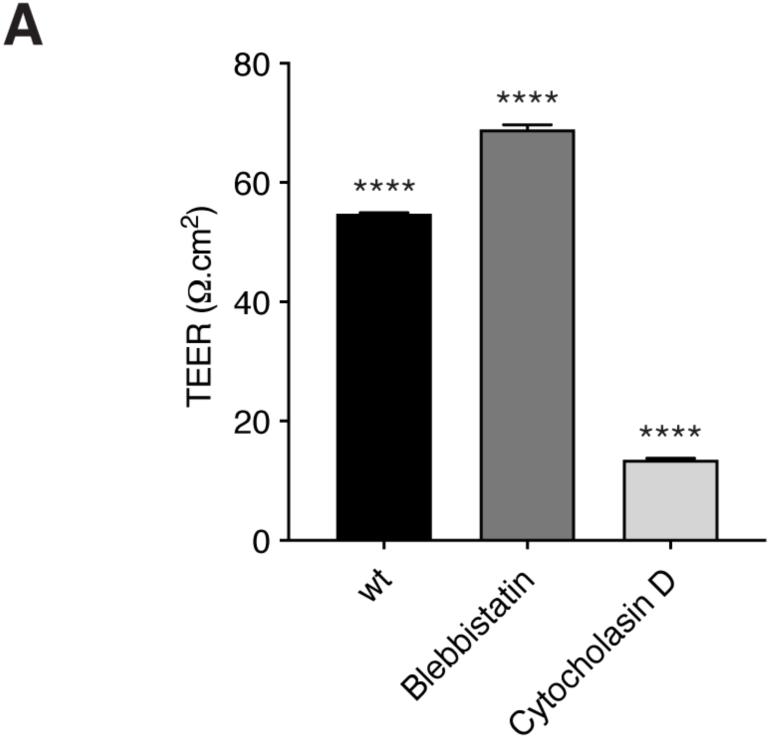
Actin contractility is detrimental to epithelial barrier function. (A) TEER measurements of wt MDCK II cells treated with DMSO, 100 μM blebbistatin, or 10 μM cytochalasin D for 3 hr. Measurements are from day 4 of confluency. Bars represent mean ± SEM, n=4, (p-values determined using a multiple comparison one-way ANOVA, p-values represent comparison all samples, **** p<0.0001)

**Supplemental Figure 2.**
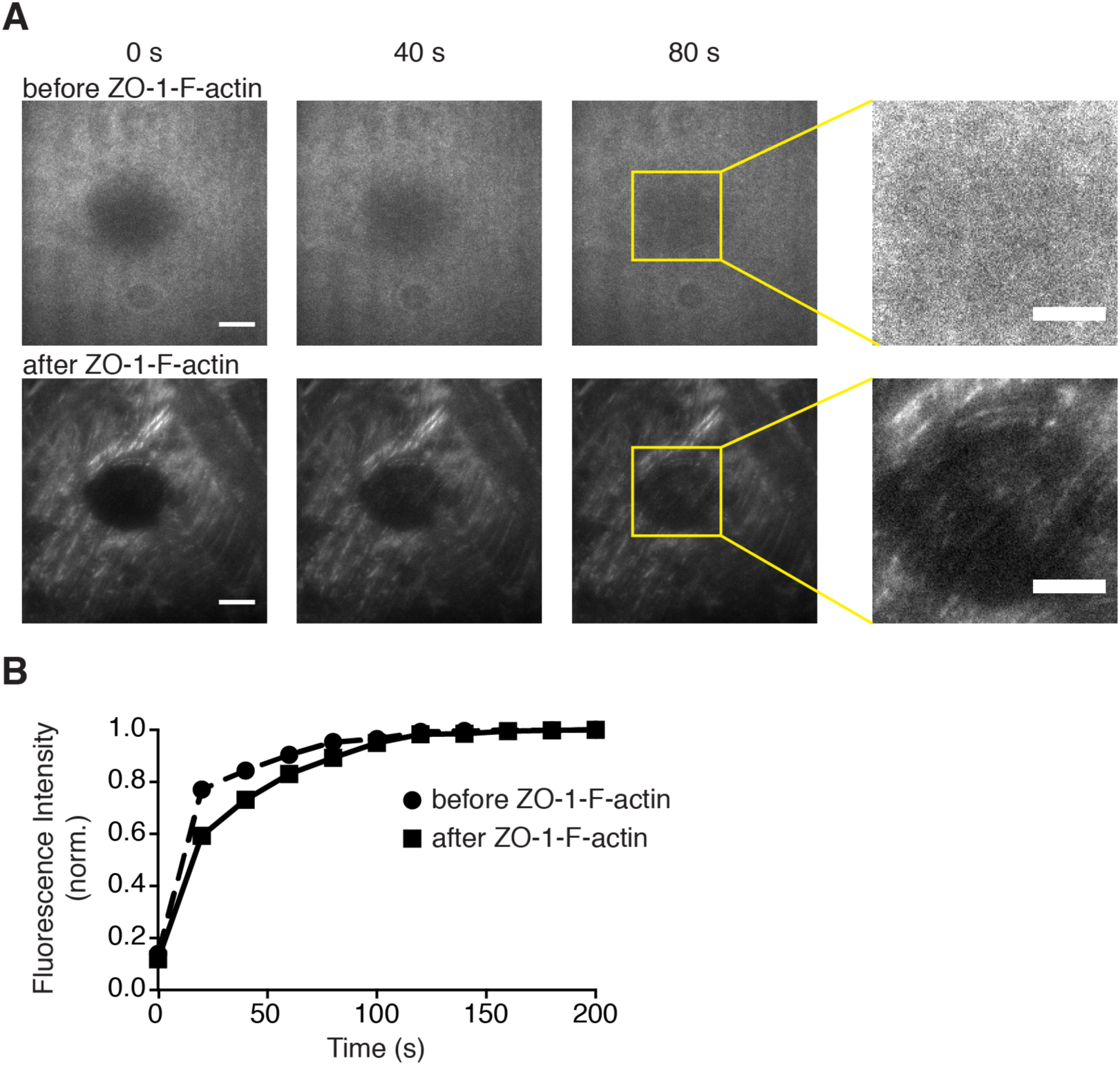
ZO-1-F-actin complexes template CtermCldn4 peptide recovery on a supported lipid bilayer. (A) Fluorescent micrographs of CtermCldn4 peptide recovery after photobleaching on an SLB with and without ZO-1-F-actin complexes. CtermCldn4 recovery is heterogeneously templated along ZO-1-F-actin complexes in the bleached area (right). Scale bar, 20 μm (left) and 5 μm (right). (B) Quantification of CtermCldn4 fluorescence intensity in bleached region over time.

**Supplemental Figure 3.**
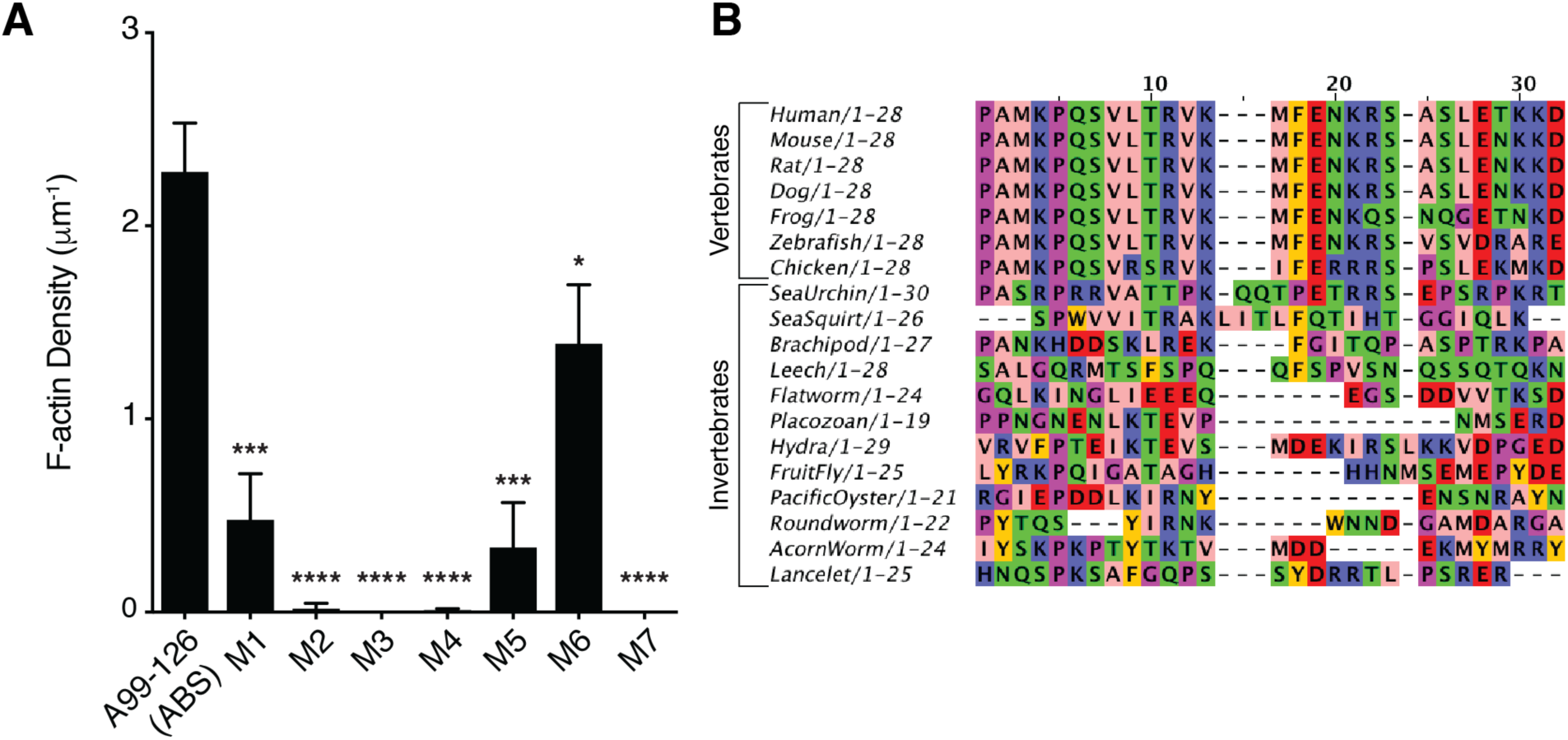
ABS alignment and mutational analysis. (A) Quantification of F-actin binding assay for ABS mutations. M1-M7 represent sequences of ABS where a string of four successive amino acids are mutated to alanine residues. Bars represent mean ± SD, n=3, (p-values determined using a two-sample t-test with A99-126, * p<0.05, *** p<0.001, **** p<0.0001). (B) ABS sequence alignment across metazoans. Sequence homology is high for vertebrates.

**Supplemental Figure 4.**
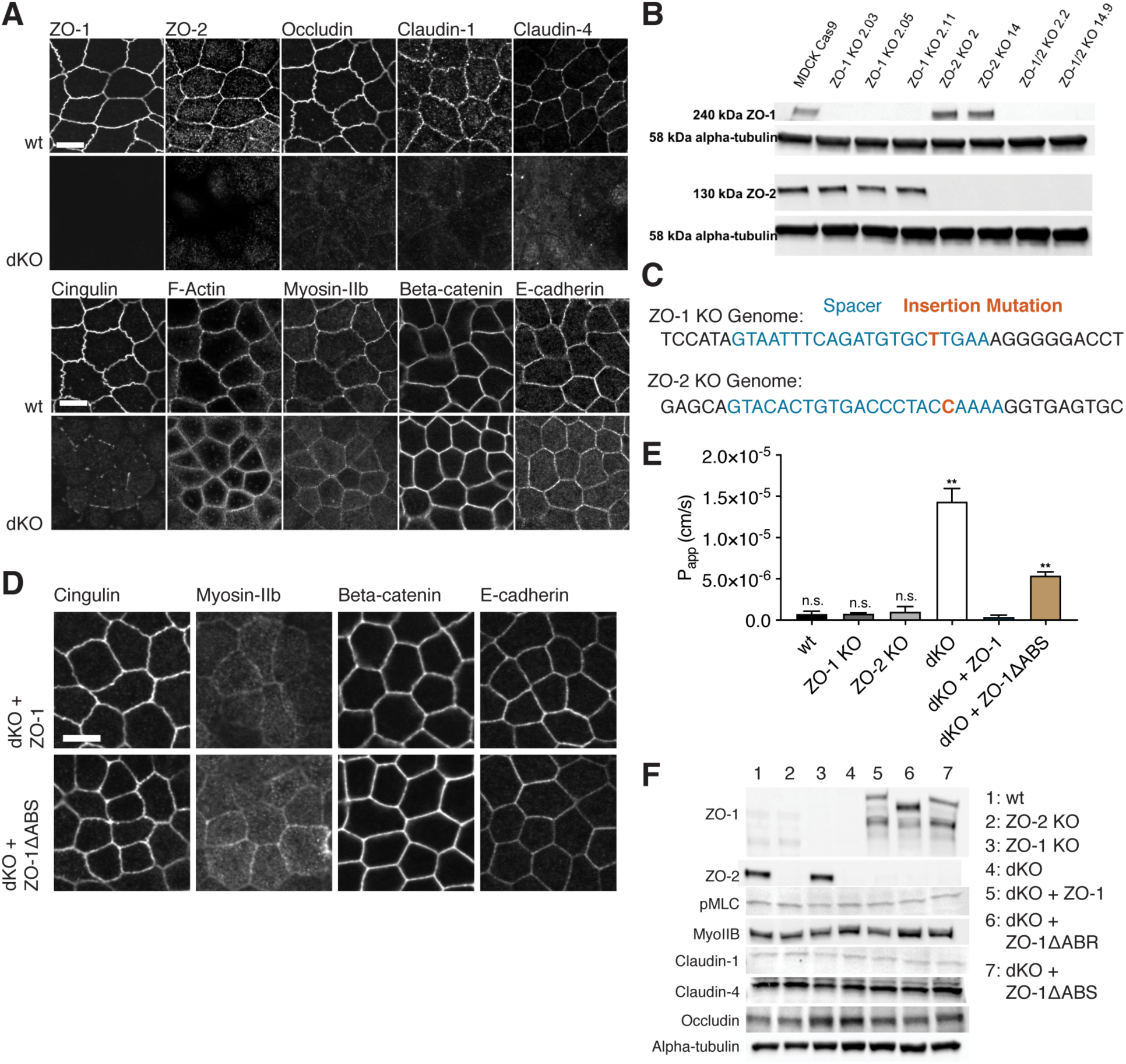
ZO-1ΔABS expression restores TJ protein localization but not barrier function compared to wt ZO-1 in dKO cells. (A) Immunofluorescent micrographs of TJ and AJ proteins in dKO and wt cells. Scale bar, 10 μm. (B) Western blots of wt and KO clones showing knockout of ZO proteins. (C) Genomic sequences of ZO-1 and ZO-2 loci in dKO cells. (D) Immunofluorescent micrographs of junctional proteins in dKO cells expressing either ZO-1 or ZO-1ΔABS. Scale bar, 10 μm. (E) Apparent permeability (P_app_) measurements of wt, KO, dKO and dKO cells expressing ZO-1 and ZO-1ΔABS are consistent with TEER results. Bars represent mean ± SEM, n=3, (p-values determined using a two-sample t-test comparison with dKO + ZO-1, ** p<0.01, n.s. p>0.05). (F) Western blot analysis of wt, ZO-1 KO, ZO-2 KO, dKO, and dKO cells expressing ZO-1 and ZO-1ΔABS.

**Supplemental Figure 5.**
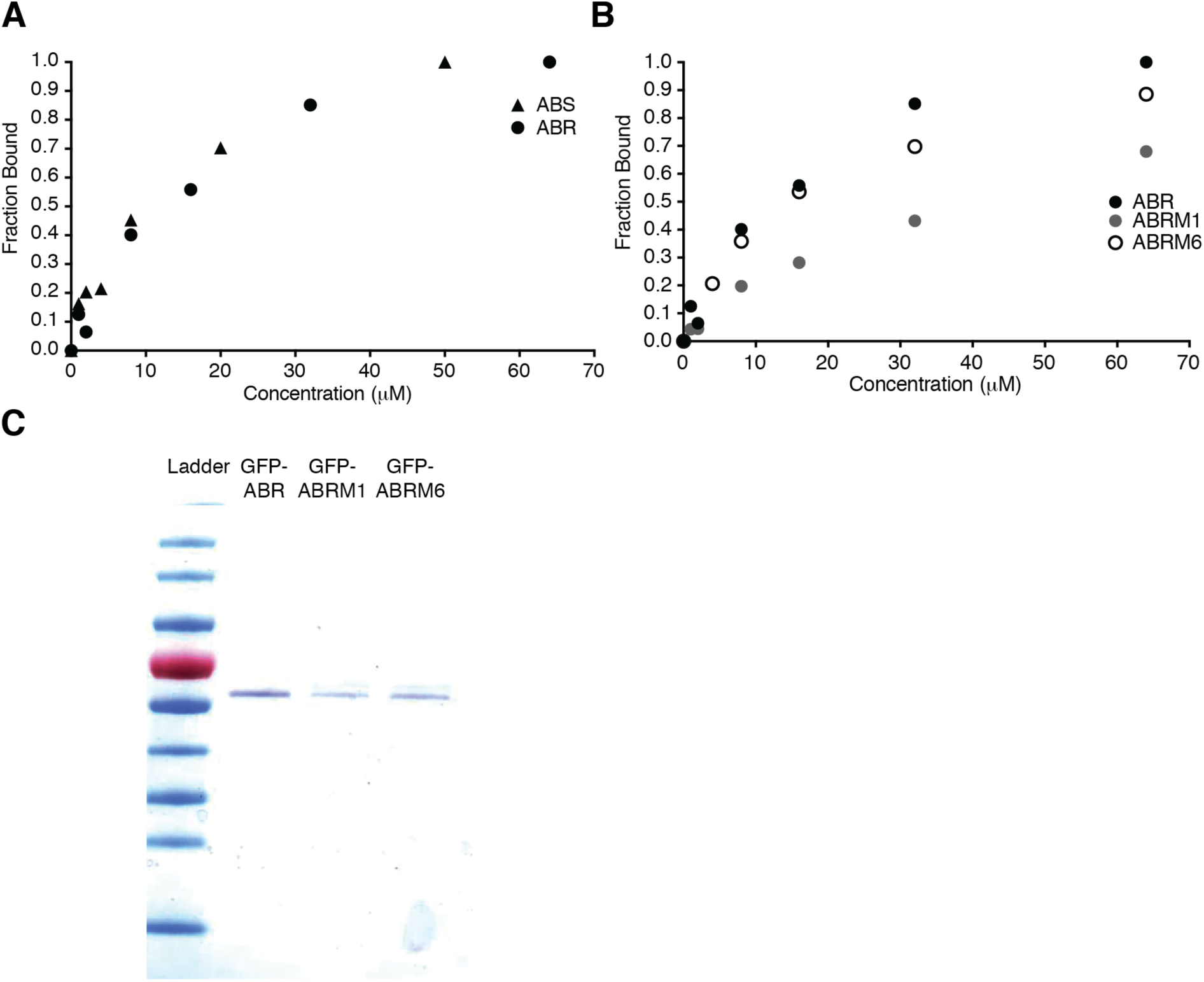
Binding curves of ZO-1’s ABS and ABR toward F-actin. (A) F-actin binding curves for ZO-1’s ABS and ABR. Both show similar affinity for F-actin. (B) F-actin binding curves for ZO-1’s ABR (K_D_ = 10.2 μM) and the ABR mutants, M1 (K_D_ = 36.7 μM) and M6 (K_D_ = 13.7 μM) (see SI Fig. 3 for more details). (C) SDS-PAGE gel of purified ZO-1 constructs: ABR, ABRM1, and ABRM6.

**Supplemental Figure 6.**
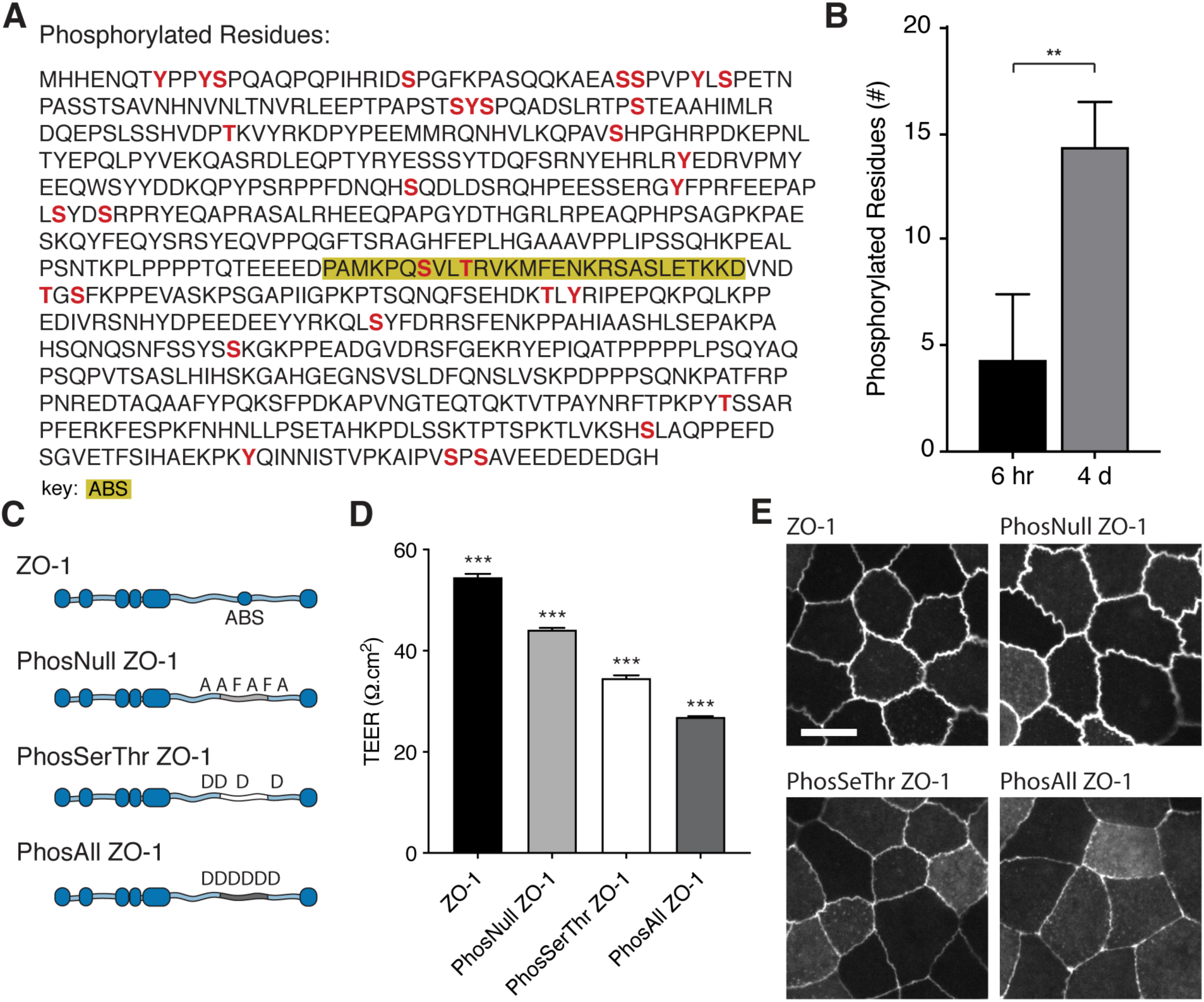
Phosphorylation of ZO-1’s disordered region varies during cell polarization and alters barrier function. (A) Sequence of ZO-1’s disordered region. Phosphorylated residues identified by mass spectrometry are highlighted in red. The ABS is outlined in gold. (B) Number of phosphorylated residues within ZO-1’s disordered region after 6 hr or 4 days of cell culture. Bars represent mean ± SEM, n=3, (p-values determined using a two-sample t-test, ** p<0.01). (C) Schematic of ZO-1 phosphorylation mutant constructs introduced into dKO cells. (D) TEER measurements of dKO cells expressing ZO-1 phosphorylation mutant constructs. Measurements are from day 4 of confluency. Bars represent mean ± SEM, n=3, (p-values determined using a multi-comparison one-way ANOVA between all conditions, *** p<0.001). (E) Fluorescent micrographs of dKO cells expressing ZO-1 phosphorylation mutant constructs. All ZO-1 mutants localize to the TJ. Scale bar, 10 μm.

## SUPPLEMENTARY TABLES

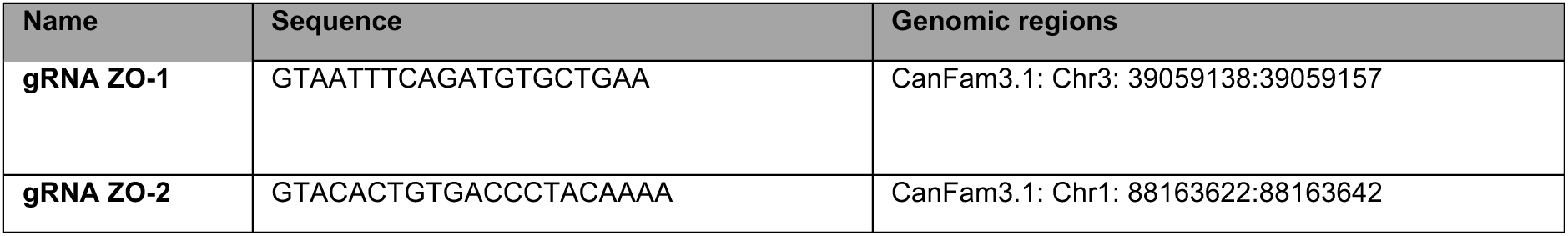

**Table S1.**
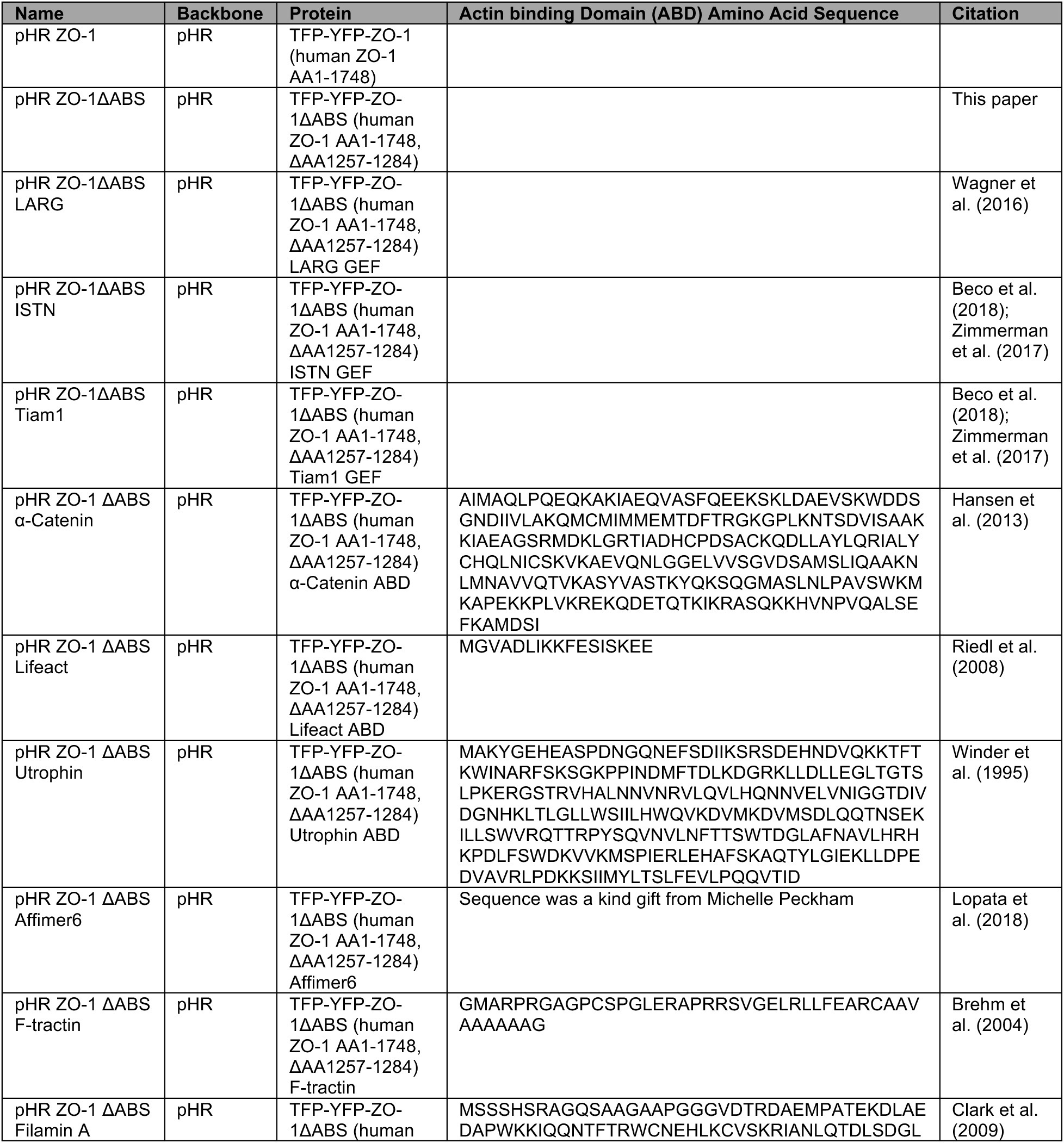
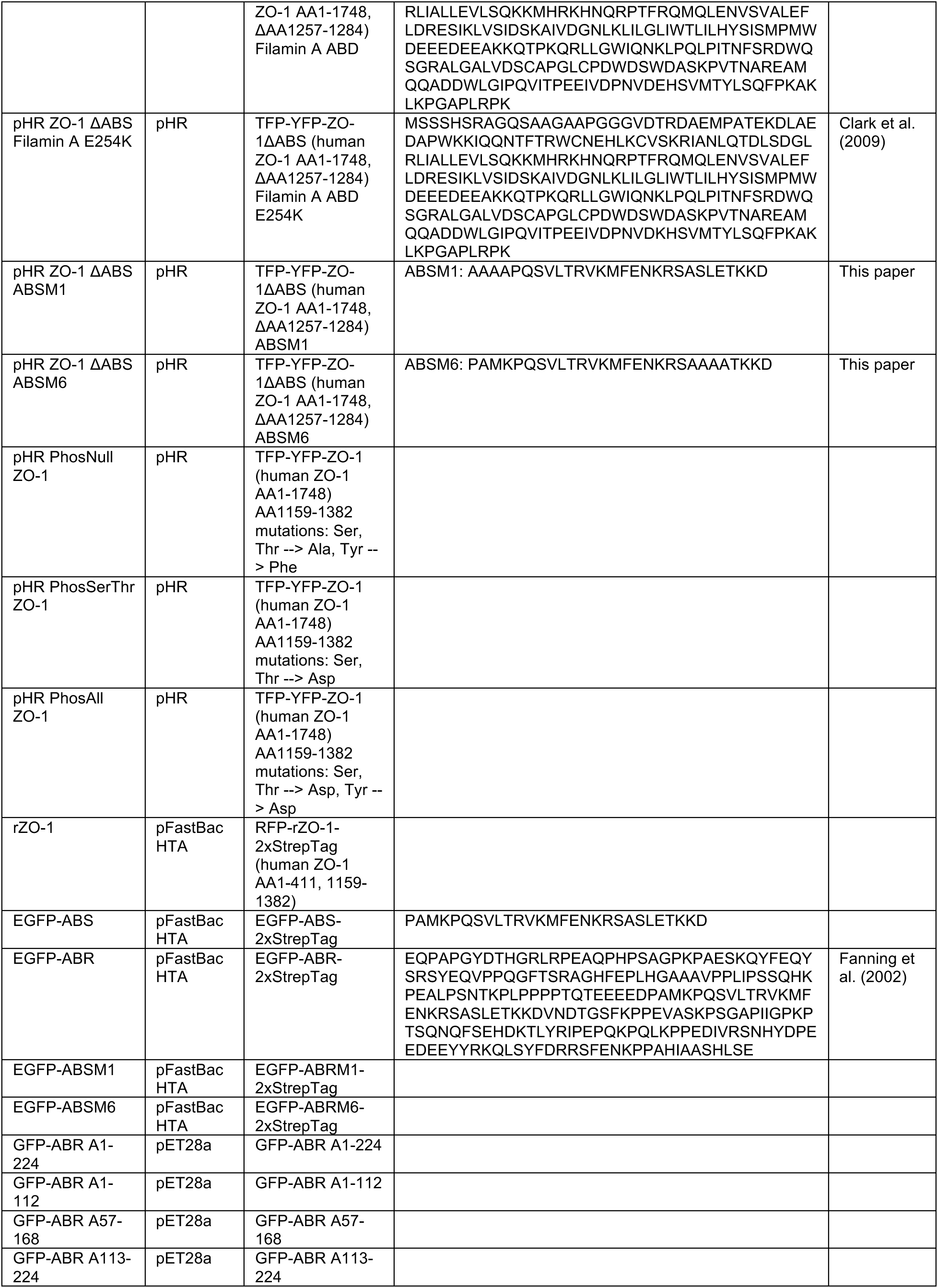
Guide RNA (gRNA) used for CRISPR/Cas9 knockout of ZO-1 and ZO-2

**Table S2.**
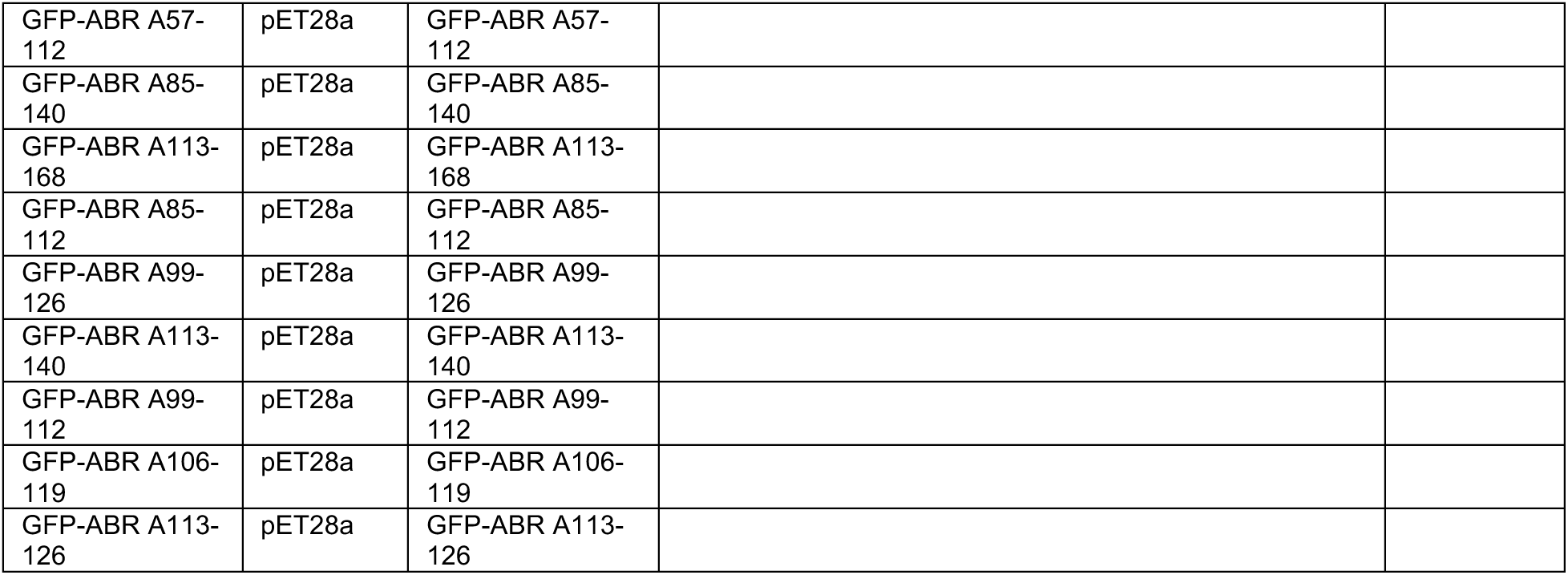
Plasmids used with amino acid sequences of actin-binding domains.

